# Environmental complexity reveals memory-guided search as the locus of learning in prey capture

**DOI:** 10.64898/2026.06.28.735138

**Authors:** Aidan M. Schneider, James N. McGregor, Mark Song, Jacob L. Amme, Samantha Zheng, Doran Wu, Jianhong Tu, Gavin Yao, Emmalee Eslinger, Jay Chitalia, Jack Powers, Varun Sinha, Eva L. Dyer, Daniel Levenstein, Keith B. Hengen

## Abstract

Ethological behavior requires the integration of perception, memory, and decision-making, yet standard laboratory tasks are often too simplified to recruit these processes jointly. Here, mice that were expert hunters in a bare arena were challenged to capture live insect prey among objects that occlude vision and afford opportunities to hide. This revealed an axis of learning absent in an empty arena: mice improved not by refining pursuit but by reorganizing how they searched for concealed prey. An agent-based model showed that short-term memory and learning of object-specific values are sufficient to account for the behavior demonstrated by mice. A compact ethogram of object-directed actions revealed that animals selectively acquired the interactions most likely to expose prey. These results show that environmental complexity recruits memory- and value-guided search strategies. Reproducible with inexpensive materials, the paradigm and its open-source analyses offer a sensitive read-out of cognitive processes largely unobservable in conventional assays.

## INTRODUCTION

Natural behaviors engage multiple neural systems operating jointly in the ecological contexts in which they evolved^1–6^. So-called ethological tasks therefore offer a powerful opportunity to study perception, movement, learning, and decision-making as integrated processes, rather than as isolated computations. In contrast to highly controlled laboratory paradigms, such as head-fixed stimulus discrimination, naturalistic behaviors unfold through continuous interaction with the environment, requiring animals to flexibly adapt their strategies in response to uncertainty and constraint. Despite this promise, ethological behaviors are increasingly studied under experimentally controlled laboratory conditions that often sacrifice the richness of natural environments in order to reduce variability. In contrast, analyses of animal behavior in the field seek to describe behavior in highly variable environments, but as a result are often restricted to simplified, coarsely defined behavioral categories. For example, studies may focus on courtship displays rather than mate-search strategies^7–10^, territorial encounters rather than patrol or exploration dynamics^11^, or consummatory actions rather than the behaviors that precede them^9,12^. Consequently, key questions remain poorly resolved: how do animals search for sparse and unpredictable resources, how do animals adapt strategies over time in the face of environmental complexity, and how does learning unfold differently in naturalistic, unconstrained tasks relative to their sparsified laboratory versions?

Prey capture is an ideal example of an increasingly well-studied ethological paradigm^1,13–21^ that remains limited in important ways by experimental design decisions. In laboratory studies, prey capture is often examined in empty or weakly structured environments, and even when environmental richness is introduced, interactions with the environment are rarely analyzed directly, despite the fact that the vast majority of animal behavior is impossible to understand without knowledge of the environment, broadly defined. When environmental factors are considered at all, their role is inferred through effects on capture probability, latency, approach characteristics, or efficiency^1,21–23^. In contrast, field studies frequently focus on narrow behavioral moments, such as the instant of capture, with limited access to the prior behavioral sequence comprising search and pursuit^24–27^. Prior work has nonetheless been highly productive within these constraints, focusing primarily on the biomechanics, sensorimotor control, and motor coordination that mechanistically account for the acute aspects of prey capture^1,13–21^. However, for many species with the requisite cognitive capacity, prey capture in nature presumably involves the integration of sensory processing, locomotion, valuation, and decision-making, and unfolds across distinct behavioral phases spanning multiple orders of magnitude from milliseconds to tens of minutes. As in many areas of ethology, these cognitive and strategic dimensions are often neglected in favor of summary descriptions (like success/failure) or biomechanical characterizations.

A second, more subtle limitation lies not in how prey capture is studied but in how it is measured, since the metrics most commonly used to characterize it are poorly suited to capturing complex cognitive features such as behavioral strategy. These metrics are intrinsic to task outcome: capture probability, time to capture, or predator–prey distance^1,14,16,21^. While informative, these metrics primarily index improvements in performance and offer limited insight into how the behavior is organized on multiple scales. An animal may, for example, exhibit similar capture times across trials while fundamentally altering *how* it searches the environment or allocates effort across behavioral states. In parallel, an emerging paradigm in behavioral analysis uses unsupervised segmentation methods to identify recurring behavioral modules^28–33^ without reference to task structure or environmental context. Although powerful for discovering patterns, these methods often quantify behaviors that seem indirectly related to the ethological process of interest. While such approaches can in theory be carried out in enriched environments (if challenges with animal tracking and event timestamping are overcome), environmental structure is not explicitly leveraged, leaving behavior–environment coupling difficult to interpret mechanistically.

Understanding prey capture as an ethological process requires a framework that connects behavioral structure to the environment that shapes it. Summary statistics collapse this coupling into outcome measures, while the unsupervised approaches described above leave it implicit, making behavior–environment relationships difficult to interpret mechanistically. A minimal, interpretable, agent-based model offers a way forward: by specifying the small set of constraints sufficient to produce structured behavior, it makes explicit how that structure may arise^34,35^. Formalizing only the components required for the process to unfold—visibility, pursuit, hiding, and search—links qualitative and quantitative features of behavior directly to task structure rather than absorbing them into unexamined parameters. The aim of such an approach is to identify the minimal mechanisms that give rise to adaptive organization in complex environments.

Here, we study prey capture in cluttered environments where objects shape visibility, movement, and learned value. We show that search is the dominant and most flexible component of the task, and that learning reorganizes search across timescales more than it refines pursuit. Using unsupervised analyses of movement trajectories, we identify the principal sources of behavioral structure in our data. We then introduce a minimal agent-based predator–prey model that reproduces key features of the observed behavior, showing that these effects reflect behavior–environment interactions, emerge as the predator learns the environment, and require memory. Finally, we classify environment-referenced actions to identify the object-directed behavioral changes that accompany learning. Together, these results show how adaptive behavior in complex environments emerges through the reorganization of search shaped by environmental structure, offering a principled framework for studying learning in naturalistic tasks.

## RESULTS

### Experimental Design and Capture Performance

Six (3 ♂, 3 ♀) C57BL/6 mice (Mus musculus) were trained individually to hunt red runner cockroaches (Shelfordella lateralis) in circular hunting arenas (36.83 cm H × 25.87 cm D), which were empty aside from bedding (Fig. 1A, left), as previously described^21^. Briefly, mice were initially acclimated to three cockroaches for two consecutive nights (12 h dark periods). Following acclimation, mice were food deprived overnight prior to the sequential introduction of six roaches, each following the termination of the prior trial due to capture or time out (*>* 30 minutes without capture) (Fig. 1B). All six animals exhibited robust learning in this sparse context, and capture times were asymptotic within the first four days. Despite the early emergence of stable expert performance, trials were continued for 10 additional weeks to ensure consolidation of prey capture behavior in simple conditions.

**Figure 1:**
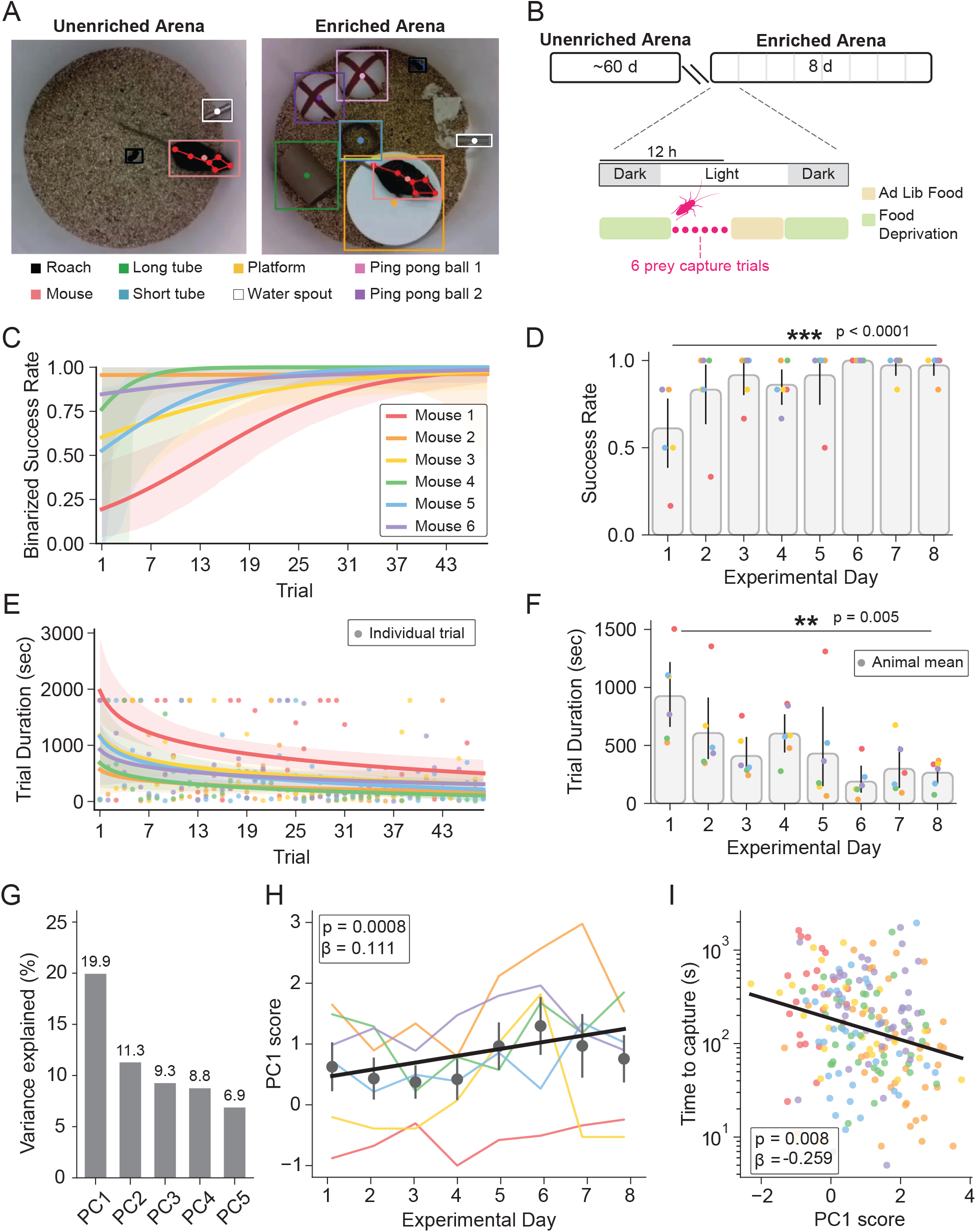
Mice learn to successfully capture prey in a cluttered environment. **A**. Overview of prey capture environments. (Left) In typical prey capture experiments, roaches are dropped into the empty, circular, home-cage chamber. (Right) We employ a novel assay in which various objects are placed throughout the arena, meant to simulate the cluttered environment in which prey capture occurs in the wild. **B**. Experimental timeline. Mice are initially trained in an empty arena for many days. After achieving expert-level performance, mice are then tested in the enriched prey capture task in a cluttered arena for eight days (Experimental Days 1-8). **C**. Logistic fits for individual animals’ success rates, where success is defined by the mouse capturing the roach within 1800 s (30 min). If the mouse fails to capture the roach within this time, the trial counts as a failure and the roach is replaced for the subsequent trial. **D**. Mean success rate on each experimental day of enriched prey capture. Each dot represents the mean success rate of each individual animal. (*** p < 0.001; linear mixed effects model: success ~ experimental day + (1|animal), where animal is included as a random effect with post hoc emmeans contrasts to compare Day 1 vs. Day 8) **E**. Logistic fits for individual animals’ trial times over the course of all trials. 1800 s is the maximum allowed trial time before considering the trial a failure. Each dot represents the duration of one individual trial. **F**. Mean trial duration on each experimental day of enriched prey capture. Each dot represents the mean trial duration for each individual animal. (** p < 0.01; linear mixed effects model: trial duration ~ experimental day + (1|animal), where animal is included as a random effect with post hoc emmeans contrasts to compare Day 1 vs. Day 8) **G**. Variance explained by the five leading principal components of the per-frame movement features. **H**. Trial-level PC1 scores across experimental days. Colored lines show per-animal daily means (colors as in C), black points show mean *±* SEM across animals, and the black line shows the model fit. (*p* = 0.0008; linear mixed effects model: PC1 ~ experimental day + (1|animal), where animal is included as a random effect.) **I**. Trial-level PC1 score versus time to capture. Each dot represents one trial, colored by animal. Higher PC1 predicts faster capture. (*p* = 0.008; linear mixed effects model: log(time to capture) ~PC1 + experimental day + log(number of tracked frames) + (1|animal))

Mice then completed an eight-day paradigm of prey capture in a complex environment (Fig. 1B). These complex hunting environments featured a set of four types of upcycled items: 1) a large PVC pipe cap (platform), 2) a bathroom tissue roll core (long tube), 3) a short segment of a bathroom tissue roll core (short tube), and 4) two separated halves of a jumbo ping-pong ball (Fig. 1A, right). Primarily, these objects served to increase environmental complexity and encourage novel interactions between the mouse and roach. Secondarily, this setup is reminiscent of the cluttered anthropogenic environments in which small rodents and insects thrive and interact (e.g., garbage containers, attics, etc.). Notably, in this particular pair of species and experimental conditions, pre-training in a less cluttered environment appears to be critical, as three naive mice whose initial exposure to roaches was overnight in the cluttered context failed to hunt.

To track the position of animals and objects during trials, we used multiple deep learning-based pose estimation models^36,37^. A convolutional model for pose estimation^36^ previously trained to track key points on a single mouse in the empty environment was employed to quantify mouse position and posture (Fig. 1A). Occasional occlusions introduced by objects in the arena (e.g., when mice moved through the long tube) were sufficiently infrequent that undetected keypoints were treated as missing and ignored. A convolutional model for object detection^37–39^ tracked the roach’s position within an estimated bounding box. Due to its small size relative to the objects, the roach was more frequently occluded, often hidden from view for tens of seconds. To overcome this, we relied on an annotation-heavy strategy where a human scorer labeled the roach’s position in one of every 15 frames (i.e., one labeled frame per second). Manual labels were used to supplement and constrain model predictions. A second model for object detection was trained to continuously label the position of the listed inanimate objects with a bounding box.

Over the course of days 1-8, hunting success in the complex environment increased markedly, converging to near 100% success across animals on later days (Fig. 1C, D) (*p* < 0.0001; linear mixed effects model: success ~ experimental day + (1|animal), where animal is included as a random effect with post hoc emmeans contrasts to compare Day 1 vs. Day 8). Intuitively, trial durations were consistently much longer and more variable in the complex environment than the empty arena. Despite this, trial durations decreased significantly over time (Fig. 1E, F) (*p* = 0.005; lmer: trial duration ~ experimental day + (1|animal)). Trials that lasted 30 minutes were terminated by the experimenter and categorized as failures. Failures were a large contributor to trial duration variance, although significance between Day 1 and Day 8 was maintained even when excluding failures from analyses. As an initial, unsupervised survey of how behavior changed over learning, we performed principal component analysis on per-frame movement features, where the leading component explained 19.9% of the variance (Fig. 1G). Trial-level scores on the leading component (PC1) increased systematically across experimental days (*β* = 0.111 per day, *p* = 0.0008; Fig. 1H), and higher PC1 predicted faster capture at the individual trial level, controlling for experimental day and number of frames (*β* = −0.259, *p* = 0.008; Fig. 1I). Unsupervised dimensionality reduction of the same features qualitatively recapitulated this structure, revealing prominent clusters that capture trial progression and individual variability across animals (Supplementary Fig. S1A,B). Taken together, these results demonstrate that mice with extensive empty-arena experience still adapt behaviorally and improve prey capture performance when confronted with complex environments.

### Behavioral states and kinematics

Consistent with our hypothesis that environmental enrichment should promote adaptive alterations in murine predatory strategies, prey capture dynamics shifted categorically after mice transitioned from the empty environment to the complex, cluttered chamber. Hunting in the empty environment was typified by a single short, ballistic pursuit. This method was effectively obviated in the cluttered environment as roaches often hid for tens of seconds. While mice could, in principle, simply chase roaches if/when they emerged from hiding, mice instead reliably exhibited a search-based hunting strategy. Searches were often successful, resulting in the mouse re-initiating pursuit. Such cyclic alternation between search and ballistic pursuit typically spanned minutes prior to capture and consumption. Despite the added challenge of search, mice were persistent and rarely appeared to lose interest in hunting.

We segmented trials into three states: pursuit, search, and abandonment. A panel of three independent human scorers achieved *>*90% pairwise agreement across all trials. As broadly defined states, pursuit and search were trivially divisible based on simple behavioral metrics such as speed and total displacement. Across all trials, mouse locomotor speed was significantly higher during pursuit than during search (and much higher than abandonment) (Fig. 2A; *p* < 0.0001, lmer: speed ~ state + (1|animal)). Additionally, mouse locomotor speed was highly correlated with the roach locomotor speed during pursuit, whereas the two were uncorrelated during search (Fig. 2B; *p* < 0.0001, lmer: correlation ~ state + (1|animal)). Further, consistent with the category labels, mice oriented their bodies towards the roach during pursuit but not search (Fig. 2C,D).

**Figure 2:**
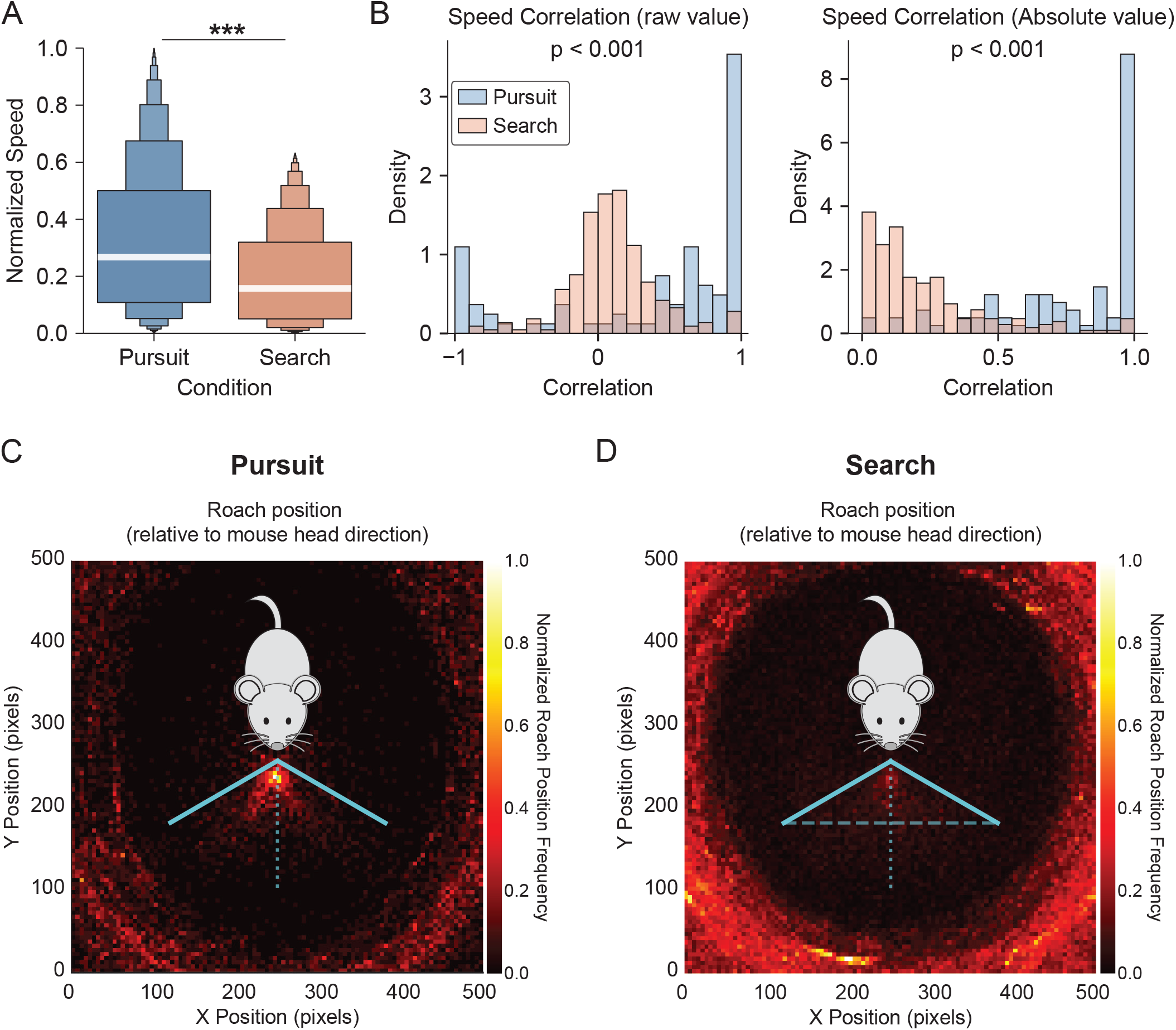
Mice reliably modify their movement speed and body orientation during enriched prey capture trials. **A**. Mouse speed, pooled across all trials and all mice and normalized to the maximum value, during pursuit and search states. Mouse speed is significantly higher during pursuit than during search (*p* < 0.0001, linear mixed effects: speed ~ state + (1|animal) where individual animal is a random effect). **B**. Histogram of raw correlation (left) and absolute value of correlations (right) between mouse and roach speed during trials, pooled across all trials and animals. In both cases, speed was more highly correlated between mouse and roach during pursuit (*p* < 0.0001, linear mixed effects: correlation ~ state + (1|animal) where individual animal is a random effect). **C**. Heatmap of roach position, relative to mouse head direction, across all video frames. During pursuit, roaches were often located in front of the field of view of the mouse. **D**. Same as in C, but during search. In this condition, roaches were often either hidden or along the edges of the chamber.

We next examined these states as a function of hunting experience in the complex environment. Search was the predominant state (accounting for ~80% of trial time), followed by pursuit (~15– 20% of trial time), and occasional abandonment (<2% of trial time, which we were statistically underpowered to examine further) (Fig. S2A). Across the 48 trials per animal, the fraction of trial time spent in pursuit and search did not change with trial number (pursuit: *p* = 0.612; search: *p* = 0.891; lmer: fraction of time in state ~ trial + (1|animal); Fig. S2A). The number of bouts per trial was likewise stable in search (*p* = 0.0851) and showed only a marginal decrease in pursuit (*p* = 0.0491; lmer: bout count ~ trial + (1|animal); Fig. S2B). Overall, the gross temporal structure of pursuit and search—how the animal partitioned each trial between the two states—remained largely stable across learning, indicating that the behavioral changes we describe below arise within the search state rather than from a reallocation of time between states.

We quantified discrete aspects of mouse movement as a function of experience (Day 1 vs. Day 8) to test for sensorimotor learning, the primary form of learning documented in previous prey capture studies^1,13–21^. Each feature was computed over four temporal windows spanning < 100 ms to 1 s (67, 333, 667, and 1000 ms).

During pursuit, mouse kinematics were largely stable across learning. Neither body-centroid travel speed nor angular speed changed in mean at any timescale (all *p >* 0.1; Fig. S3A, B), and their variability was similarly unchanged (angular: all *p >* 0.39). Mouse head speed increased modestly, but only at the shortest timescale (67 ms: *p* = 0.026; longer timescales *p >* 0.1; Fig. S3C), with no accompanying change in its variability (all *p >* 0.24). Mice also did not orient their heads toward the prey more often with experience (*p* = 0.55). The prey’s behavior during pursuit was likewise unchanged: neither mouse–roach distance (*p* = 0.744) nor roach speed (*p* = 0.096) differed across days (Fig. S3D). Together, these results indicate that pursuit kinematics were largely fixed, with only subtle modification of the fastest head movements.

Search kinematics, by contrast, changed markedly across learning, in both mean and variability. Body-centroid travel speed increased significantly at every timescale (all *p* < 10^−4^; Fig. S4A). Mouse head speed increased (all *p* < 10^−5^; Fig. S4C) and became significantly more variable across all timescales (*p* = 0.015 to 1.8 × 10^−5^). Angular travel speed increased across all timescales (67 ms: *p* = 0.006; 333 ms: *p* = 0.0016; 667 ms: *p* = 0.0007; 1000 ms: *p* = 0.0012; Fig. S4B), with a parallel increase in its variability (all *p* < 0.012). Despite these changes, mice did not orient their heads toward the prey more often with experience (*p* = 0.52), consistent with the prey typically being out of view during search. Unlike during pursuit, mouse–roach distance showed a non-significant trend toward change across days (*p* = 0.086), while roach speed was unchanged (*p* = 0.342; Fig. S4D). The trend toward reduced mouse–roach distance raises the possibility that mice learn the behavioral patterns, and likely hiding locations, of their prey over time.

### Unsupervised Analysis

We next sought to identify the main effects present in mouse movement throughout the environment. A standard approach is unsupervised dimensionality reduction, which projects data points—here, mouse locomotor trajectories—into a low-dimensional space where similar paths lie adjacent to one another. In this context, the main effects are the parameters that drive the separation of points across the space. Interpretability is greatest when the observable variance in the low-dimensional projection maps clearly onto a well-defined feature of the data.

To apply this approach, we first defined a meaningful coordinate system for mouse locomotor paths. In an empty circular arena, a mouse’s instantaneous position can be described succinctly by a single coordinate: its distance to the nearest wall. Introducing objects to the environment requires a richer description, so we adopted a multidimensional coordinate system capturing the mouse’s distance to the wall and to each object (ping-pong balls, short tube, long tube, and platform). We then performed dimensionality reduction on locomotor paths 1–10 s in duration by computing pairwise distances between paths with dynamic time warping and projecting them into a low-dimensional space via multidimensional scaling.

For each timescale, we colored points (each corresponding to an individual path) according to a range of continuous scalar metrics describing the path, as well as categorical descriptors that contextualize the data. Categorical variables such as animal identity and date showed no clear relationship with the space.

On long timescales (~10 s), variance in the lower dimensions was shaped by the total distance the animal traveled over the interval, and thus indirectly by many of the kinematic metrics described above (e.g., the mean and variance of speed).

On intermediate timescales (~3 s), variance was instead explained by the overall shape of the path (Fig. 4). Continuous travel around the perimeter of the enclosure was a recurring motif.

Such thigmotaxis, or wall-following, is a known behavior in captive animals, particularly under fear. Fear is an unlikely explanation here, however: our mice were extensively habituated to the task and environment, and we did not observe wall-following in the absence of roaches. A more plausible account is that roaches exhibit thigmotaxis while evading pursuit in empty arenas, and mice had extensive experience hunting in empty arenas prior to the introduction of objects. Yet the motif was no more prevalent during pursuit than during search, arguing against a pursuit-driven origin.

On short timescales (~1 s), variance was explained by the object nearest the mouse (Fig. 5A). One contributing factor may be that objects induce characteristic changes in paths as mice move through, onto, or around them. A simpler explanation is that over short intervals the mouse’s location changes minimally, so its position in the reduced space largely reflects an averaged estimate of where it is. Consistent with this spatial interpretation, the object-enriched regions of the reduced space were arranged clockwise in the same order as the objects in the arena, indicating that spatial relationships between objects are broadly preserved. Coverage of the space was not uniform, however: objects occupied regions that were not scaled to their true size. We confirmed that the proportion of time the mouse spent near each object was neither uniform across objects nor scaled to object size. Moreover, these object-time proportions were not consistently matched to the roach’s across trials. How the mouse chose to allocate time among objects was not apparent.

### Agent-Based Modeling

To guide hypotheses about how interactions between predator, prey, and environment reorganize prey capture behavior in complex environments, we constructed a minimal, mechanistically-interpretable, agent-based predator–prey model that reproduces the essential structure of the task.

The model includes two agents, predator and prey, moving within a discretized representation of the arena in fixed timesteps (Fig. 3A, top). At each timestep, each agent selects from a set of possible movements. The predator’s behavioral state is set by the visibility of the prey: when the prey lies within the predator’s visual field, the predator enters a pursuit state in which it attempts to minimize the Chebyshev distance to the prey. When the prey is not visible, the predator instead enters a search state, exploring the arena to relocate the prey. The prey follows a complementary policy: when it detects the predator within its omnidirectional field of view, it attempts to increase its distance from the predator. When it does not detect the predator, it follows a biased random policy that favors remaining still. This formulation results in straightforward pursuit of prey (Fig. 3B, top) and captures key aspects of expert-level prey capture performance in the empty arena, including near-certain success rates and relatively short capture times (Fig. 3C).

**Figure 3:**
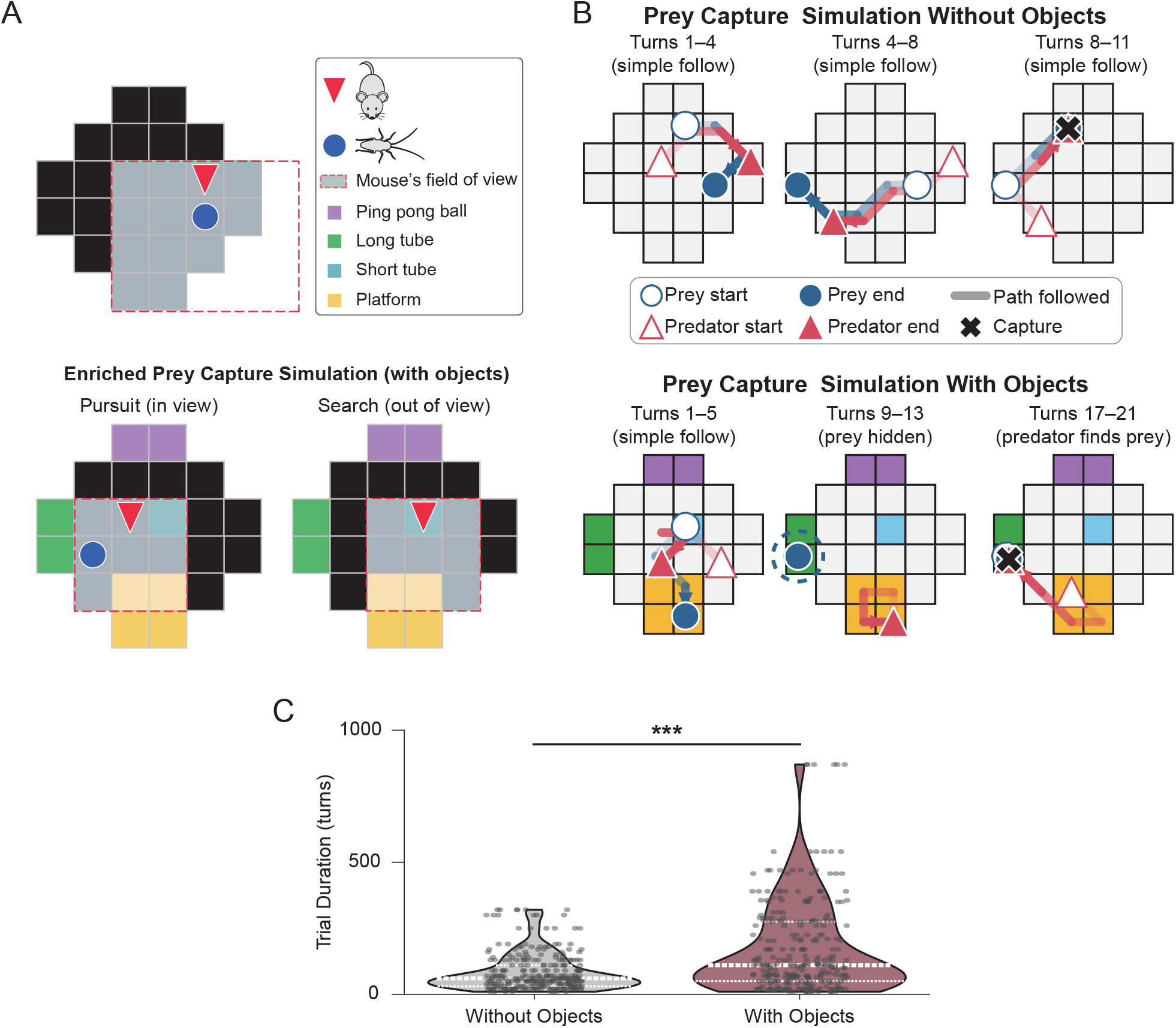
Agent-based predator-prey model reproduces key features of the prey capture task. **A**. Schematic of the agent-based model. (Top) predator (red triangle) and prey (blue circle) move through the arena in discrete timesteps; the predator’s field of view (red dashed outline) determines whether it pursues the visible prey or searches. (Bottom) With objects added, the prey can hide, requiring the predator to search to relocate it. **B**. Example simulated trials shown as a sequence of time windows (open symbols, interval start; filled symbols, interval end), for the arena without objects (top) and with objects (bottom). Without objects, the predator directly pursues and captures the prey; with objects, the prey hides (dashed ring in middle panel) and the predator must search before recapture. **C**. Trial duration (in simulation turns) for the model with and without objects, pooled across 6 independent runs of 48 trials each. Violin plots show the distribution of raw durations; dashed and dotted lines mark the median and quartiles. Trials with objects lasted significantly longer (linear mixed-effects model, log(turns) ~ group + (1|run); 1.66-fold increase, p = 0.0001).

To extend this framework to the enriched arena, we added objects that allow the prey to hide, requiring the predator to search when the prey is concealed (Fig. 3A, bottom). Object scale and layout approximated the experimental arena. Pursuit behavior was largely unaffected, as the predator re-initiated a direct chase once the prey became visible. As a result, the inclusion of objects altered predator–prey dynamics primarily during search. This minimal addition reproduced several features of the enriched-arena data, including frequent pursuit–search transitions (Fig. 3B, bottom) and substantially longer capture times than in the empty arena (Fig. 3C). To match the experimental cohort of six animals, we ran the model as six independent simulations (one per random seed), treating each run as a replicate analogous to an individual animal.

### Short-term memory of recently searched objects produces efficient, non-backtracking paths

In the agent-based model, search driven solely by random exploration caused the predator to revisit the same locations repeatedly, producing redundant, inefficient paths. We reasoned that in complex environments, prey capture may be aided by short-term memory of recently inspected objects that failed to reveal the roach, biasing search toward new objects more likely to conceal the prey (Fig. 4A). To test this, we added a short-term memory mechanism to the predator in the model that temporarily suppressed the revisitation of an empty object for a number of steps. This generated structured, non-repetitive, roughly circular search sequences that closely resembled the experimental data. Specifically, mouse trajectories on intermediate timescales (3-5 s) contained a high prevalence of semi-circular, ellipse-shaped path segments (Fig. 4B, C).

**Figure 4:**
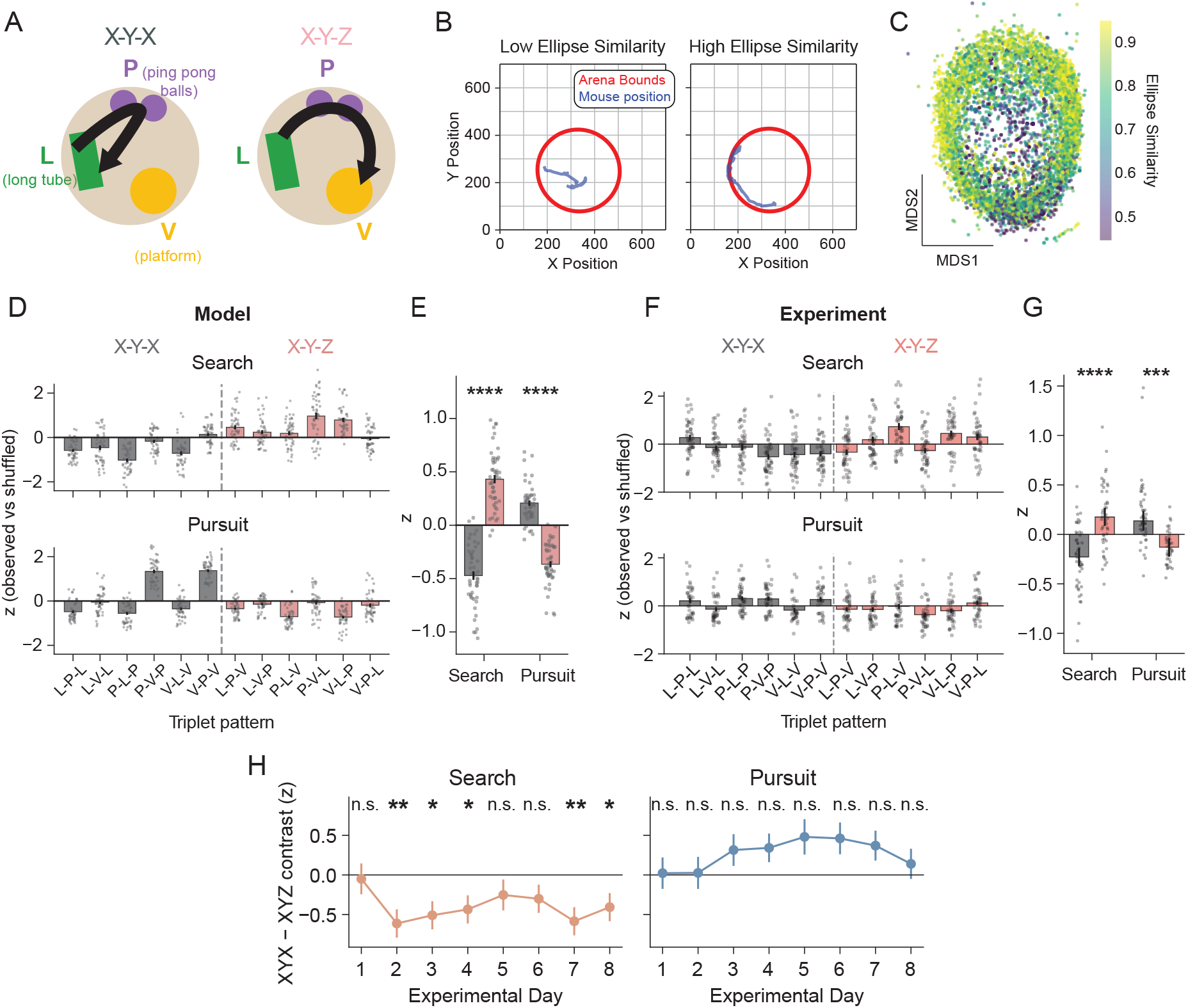
A memory-dependent search strategy biases mice toward non-repeating object sequences. **A**. Hypothesized, memory-dependent search strategy. Schematic of two possible triplet paths between objects in the arena (long tube, L; ping-pong balls, P; platform, V). Left: an X-Y-X path, in which the mouse returns to a recently visited object. Right: an X-Y-Z path, in which the mouse traverses three distinct objects. X-Y-Z paths are predicted by a memory-dependent search strategy in which the mouse avoids backtracking to recently visited locations. **B**. Two example mouse trajectories over a 5 s window. Left: low ellipse similarity, indicating a non-circular path. Right: high ellipse similarity (score *>* 0.9), indicating a semi-circular path consistent with traversal of distinct objects. **C**. Multidimensional scaling embedding of ellipse-similarity scores from 3 s segments of mouse trajectories, colored by ellipse similarity, showing a high prevalence of circular path segments across trials. **D**. Per-triplet *z*-scores (observed probability vs. null, consisting of shuffled object sequences) from the agent-based model of memory-dependent search, during Search (top) and Pursuit (bottom) states. Each dot displays the estimate from one simulation block of 6 consecutive trials; bars show mean *±* SEM across the 48 blocks (6 runs × 8 blocks). **E**. X-Y-X vs. X-Y-Z class contrast for the model, summarizing D. Marginal means from a linear mixed-effects model (z ~ triplet class × state + (1|run)) showing the X-Y-X vs. X-Y-Z class contrast by state. The model was significantly biased toward X-Y-Z paths during Search (*p* < 0.0001) and toward X-Y-X paths during Pursuit (*p* < 0.0001). Bars show mean *±* SEM. **F**. Same as D, for experimental data: per-triplet *z*-scores from mouse trajectories. **G**. Same as E, but for experimental data. Linear mixed-effects model: z ~ triplet class × state + (1|animal/day). Mice were significantly biased toward X-Y-Z paths during Search (*p* < 0.0001) and toward X-Y-X paths during Pursuit (*p* = 0.0001). *n* = 6 animals across 8 training days. **H**. Day-by-day estimated marginal means of the X-Y-X − X-Y-Z contrast for each state (error bars: SEM). Asterisks denote contrasts significantly different from zero after Benjamini-Hochberg correction within each state. The Search bias was absent on Day 1 (*p* = 0.80) and present on most subsequent days (Days 2, 3, 4, 7, 8; *p* < 0.05); no individual day reached significance for Pursuit after correcting for multiple comparisons (minimum *p* = 0.12).

To test whether such circular navigational structure reflects a memory-dependent avoidance of recently visited objects, we examined the sequences of objects mice traversed. We classified each three-object sequence (triplet) as either X-Y-X, in which the mouse returns to a recently visited object, or X-Y-Z, in which it visits three distinct objects (Fig. 4A); a memory-dependent strategy predicts an over-representation of non-repeating X-Y-Z paths during search. We first confirmed that the agent-based model produced this signature: relative to a shuffled null, the model was biased toward X-Y-Z triplets during search and toward X-Y-X triplets during pursuit (Fig. 4D,E). This split was likewise robust across the hyperparameter sweep: the model consistently favored looping trajectories during search and direct back-and-forth trajectories during pursuit across the parameter space (Fig. S6). Experimental data reproduced this pattern closely (Fig. 4F,G): mice were significantly biased toward X-Y-Z paths during search (*p* < 0.0001) and, conversely, toward X-Y-X paths during pursuit (*p* = 0.0001) (lmer: z (bias) ~ triplet class × state + (1|animal) + (1 |animal-day)). Thus, during search—but not pursuit—mice avoided backtracking to recently visited objects, exactly as predicted by a memory-dependent inspection strategy. Notably, this search bias emerged with experience: The X-Y-Z bias was absent on the first day of object exposure (Day 1, *p* = 0.80) but was significantly different from zero on the majority of subsequent days (Fig. 4H; Days 2, 3, 4, 7, and 8 *p* < 0.05, Benjamini–Hochberg corrected within state). No individual day reached significance during pursuit after correcting for multiple comparisons (minimum *p* = 0.12). This pattern suggests a learned rather than innate search strategy.

### Object-specific valuation aligns predator search with prey hiding preferences, a strategy mice learn

The agent-based model also guided a second hypothesis about strategies in complex environments: that, should a prey species be biased toward a subset of possible hiding locations, an efficient predator might learn these probabilities and subsequently concentrate its search based on this information. This factor is largely irrelevant during pursuit, when the predator remains colocalized with the prey, but becomes important once the prey hides and the predator must infer where concealment is most likely.

Mouse trajectories at 1 s resolution were organized largely by object proximity, with the nearest object explaining a substantial fraction of the variance along the first two MDS embedding dimensions (Fig. 5A). We therefore quantified the proportion of time the mice and roaches spent nearest each object over trials (Fig. 5B). To interpret these occupancy patterns, we expressed them as a predator-prey difference: for each object, the proportion of time the predator spent nearest it minus the proportion for the prey, such that values near zero indicate that the predator’s search allocation matched the prey’s distribution and large values indicate mismatch (Fig. 5C). We used the model to establish reference expectations for this measure under two search strategies. A predator that treated all objects as equally likely hiding sites (value-mismatched) distributed its inspection time independently of where the prey actually hid, producing large predator-prey differences (Fig. 5C, light bars). A predator whose search values matched the prey’s hiding preferences (value-matched) instead reallocated its inspection time to align with the prey’s distribution (Fig. 5C, dark bars; all four objects, BH-corrected *p* < 0.001), significantly reducing the total predator-prey divergence during search (Fig. 5E; *p* < 0.001). This benefit was specific to search: during pursuit, when the predator remained co-localized with visible prey, value-matching did not reduce divergence and instead significantly increased divergence (Fig. 5C,E). The realignment during search did not depend on the model’s specific parameter values: across a grid sweep over all seven prey-decision and predator search-memory parameters, a value-matched predator improved mouse–roach occupancy alignment during search relative to a value-mismatched predator (Fig. S5). These two conditions establish the range of possible predator–prey occupancy alignment and provide the templates against which we next asked whether real mice shift from the former toward the latter over learning.

**Figure 5:**
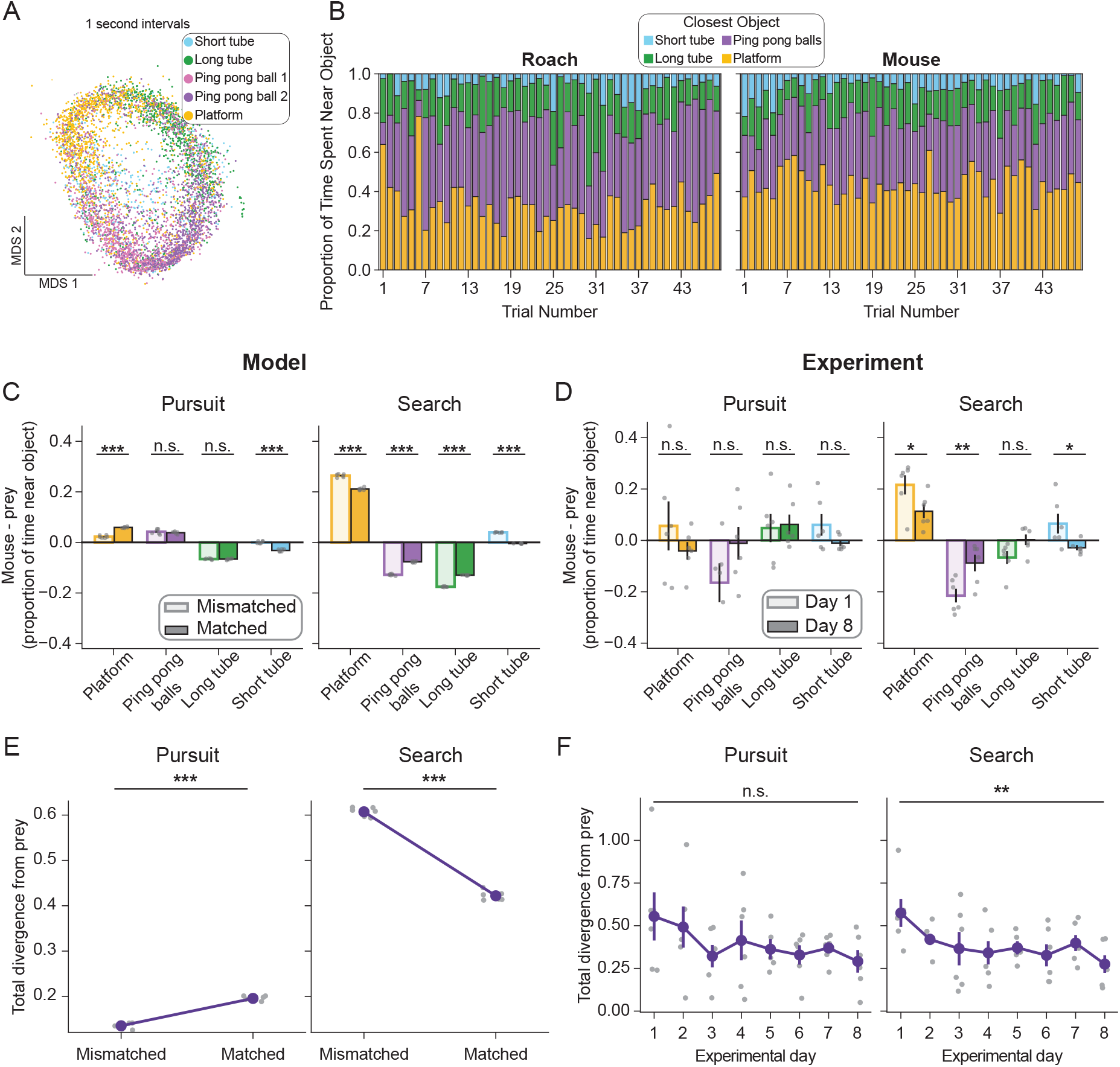
Proportion of time spent near objects reveals learned search strategies. **A**. Multi-dimensional scaling (MDS) of mouse trajectories at 1 s resolution, with each point colored by the nearest object in the arena. Proximity to individual objects explains a substantial fraction of the variance along the first two MDS dimensions. **B**. Proportion of time spent near each object on each trial, averaged across animals, for roaches (left) and mice (right). Trial number is shown on the x-axis; stacked bars sum to 1. **C**. Model predator–prey difference in proportion of time spent near each object during pursuit (left) and search (right), for value-mismatched (light) and value-matched (dark) conditions, averaged across simulated replicates and trials. Asterisks denote per-object matched vs. mismatched contrasts, BH-corrected across objects. **D**. Experimental predator–prey difference during pursuit (left) and search (right) on day 1 (light) and day 8 (dark), averaged across animals and trials. Asterisks denote per-object day 1 vs. day 8 contrasts, BH-corrected across objects. During search, the predator–prey difference moved significantly toward zero for the platform, ping-pong balls, and short tube between day 1 and day 8; no pursuit contrast was significant. **E**. Total divergence from prey (summed absolute predator–prey difference across objects) during pursuit (left) and search (right), comparing value-mismatched and value-matched conditions in the model. Bars show the mean across simulated replicates (*n* = 6); overlaid points are per-replicate means. **F**. Total divergence from prey during pursuit (left) and search (right) across experimental days. Points show the mean across animals (*n* = 6) with SEM; overlaid points are per-animal means across trials. Divergence decreased significantly across days during search but not pursuit. Statistical details and model specifications in Methods.

In experimental data, mouse object-search proportions differed substantially from the roach’s occupancy distribution during early trials but converged over learning (Fig. 5D). During search, the predator-prey difference moved significantly toward zero for the platform (*p* = 0.022), ping-pong balls (*p* = 0.006), and short tube (*p* = 0.027) between day 1 and day 8, as the mouse’s search allocation came to match where roaches most often hid; the long tube did not change significantly (*p* = 0.086). No pursuit contrast reached significance (Fig. 5D). We summarized this convergence as the total divergence between mouse and roach occupancy, which decreased significantly across days during search (*p* = 0.002) but not pursuit (*p* = 0.091; Fig. 5F). The agent-based model reproduced this pattern: relative to a value-mismatched predator, a value-matched predator showed significantly lower occupancy divergence during search (Fig. 5C,E). Mice therefore appear to learn which objects most often conceal prey and to allocate search effort accordingly.

### Classification of object-directed behaviors

Our trajectory analyses showed that objects shape how the mouse and roach move through the environment, but they do not reveal the specific behaviors mice use to engage objects during search and pursuit. To resolve these, we reduced the large space of object-related actions to a tractable ethogram. We defined three primary modes of object engagement: occupying or moving atop an object (“over”), contacting an object with the face or forepaws while adjacent (“interact”), and inspecting its interior or perimeter (“investigate”, or “through” for the long tube) (Fig. 6A). We annotated these behaviors for the ping-pong balls, long tube, and platform (the short tube was excluded owing to minimal mouse proximity and limited interactions).

**Figure 6:**
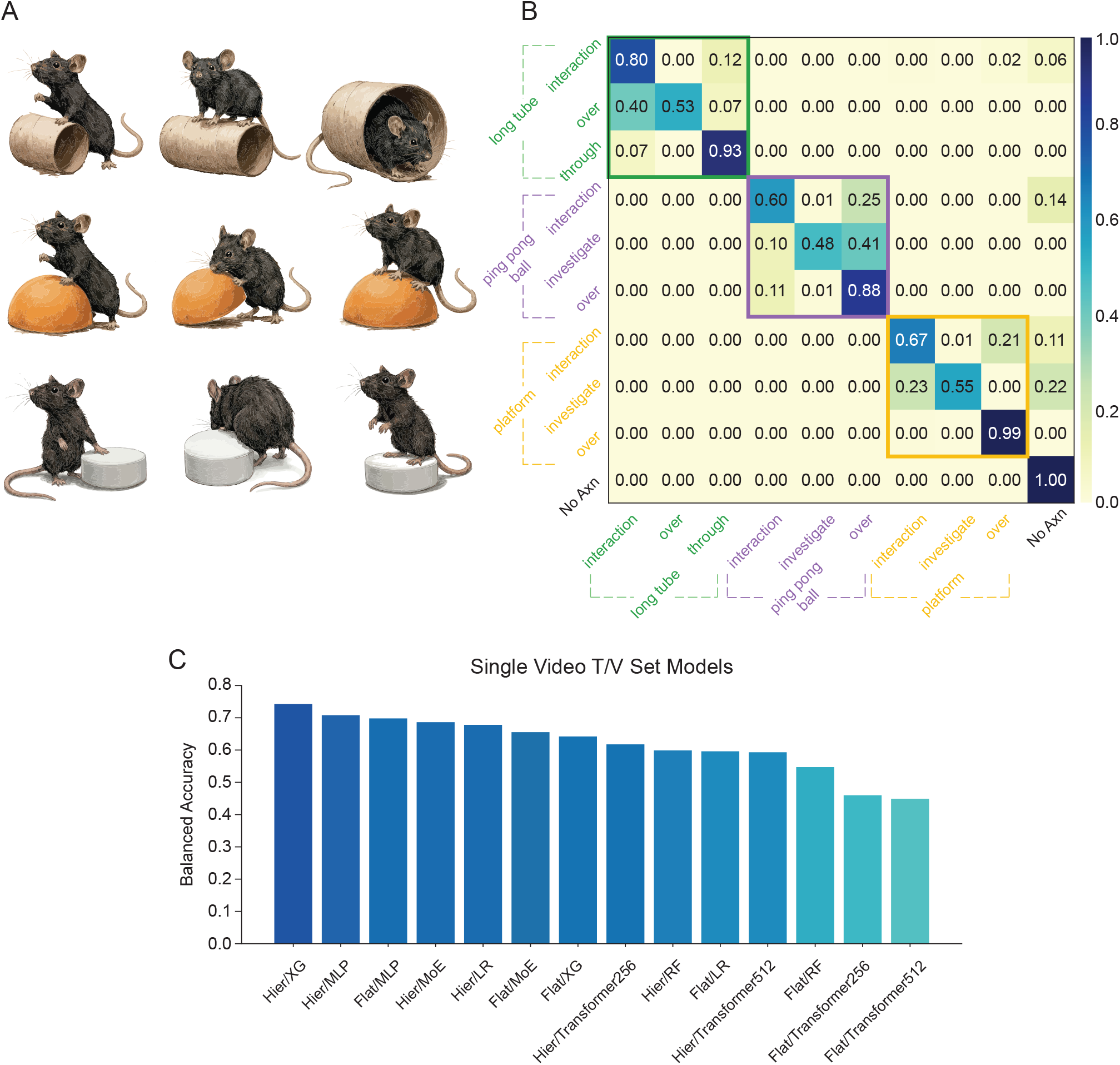
Machine learning classifier accurately labels mouse ethogram during enriched prey capture. **A**. Schematics of object-directed behaviors observed in the enriched prey capture arena. Rows depict interactions with the long tube (top), ping-pong ball (middle), and platform (bottom). **B**. Confusion matrix for hierarchical XGBoost model classifying object-directed behaviors. Colored boxes group behaviors by object: long tube (green), ping-pong ball (purple), platform (yellow). Most behaviors are classified accurately along the diagonal; the most confusable categories are within-object motif distinctions (e.g., platform interaction vs. investigation). **C**. Balanced accuracy across candidate model architectures for classifying the mouse ethogram, evaluated on held-out trials within a single train/validation split. The hierarchical XGBoost model (Hier/XG) achieves the highest balanced accuracy and was used for all subsequent ethogram analyses. Models compared include hierarchical and flat XGBoost (Hier/XG, Flat/XG), random forests (Hier/RF, Flat/RF), transformers (Transformer256, Transformer512), mixture-of-experts models (Hier/MoE, Flat/MoE), multilayer perceptrons (Hier/MLP, Flat/MLP), and logistic regressions (Hier/LR, Flat/LR).

A committee of three trained scorers manually labeled classes of object engagement in a subset of data (all trials from one day per animal). We then trained a classifier to predict behaviors across all frames from DeepLabCut pose features and YOLO-based object localization^36,37,39^ (Fig. 6B). Across models ranging from linear decoders to neural networks, a hierarchical XGBoost classifier performed best (Fig. 6C). The model first classified which object (if any) the mouse engaged with, and then the specific behavior (e.g., “over”, “interact”, “investigate”) within that object group. This structure reduced classification error, with remaining discrepancies largely reflecting frame-level ambiguity rather than clear misclassification: during transitions such as climbing onto the platform, the frame marking the shift from “interact” to “over” is inherently ambiguous yet can contribute disproportionately to error. Balanced accuracy for the hierarchical XGBoost model was 74.2% for individual behaviors (chance = 10%) and rose to 94.3% when allowing for ambiguity within object categories (chance = 25%).

### Mice learn to engage in object interactions that expose hidden prey

Objects in the arena differ in whether interacting with them can expose concealed prey. Roaches can wedge themselves between the platform and the enclosure wall, where the gap is too narrow for the mouse to pursue and the platform too heavy to displace (Fig. 7A, left). Ping-pong balls, by contrast, are light enough to be moved during investigation, allowing mice to flush hidden roaches (Fig. 7A, right). Consistent with this asymmetry, roaches were frequently located near the ping-pong balls—but not the platform—at the end of successful searches (Fig. 7B). We therefore hypothesized that, over learning, mice would selectively increase the object interactions that expose prey. To test this, we used the hierarchical XGBoost classifier to label each object-directed frame by its object and behavior type (interaction, investigate, or over/through) and tested whether each proportion changed between the first and last days of object experience (binomial GLMM, emmeans contrasts Day 1 vs. Day 8, FDR-corrected across the nine behaviors; Fig. 7C–E). During search, long-tube and platform behaviors were unchanged across learning (Fig. 7C, E). In contrast, all three ping-pong-ball behaviors during search changed significantly over learning (Fig. 7D): interaction decreased (*p* = 0.005), while investigation (*p* < 0.001) and over behaviors (*p* = 0.002) increased. Thus, as mice learned, their engagement with the ping-pong balls shifted away from simple object contact and toward the exploratory investigate and over behaviors that displace the object and can flush concealed prey. Consistent with our broader finding that learning reorganizes search more than pursuit, the object-directed changes during search were not mirrored by a coherent pattern during pursuit (Fig. S7). Together, these results indicate that mice learn to favor the object interactions that expose prey, demonstrating effective, context-dependent search behaviors.

**Figure 7:**
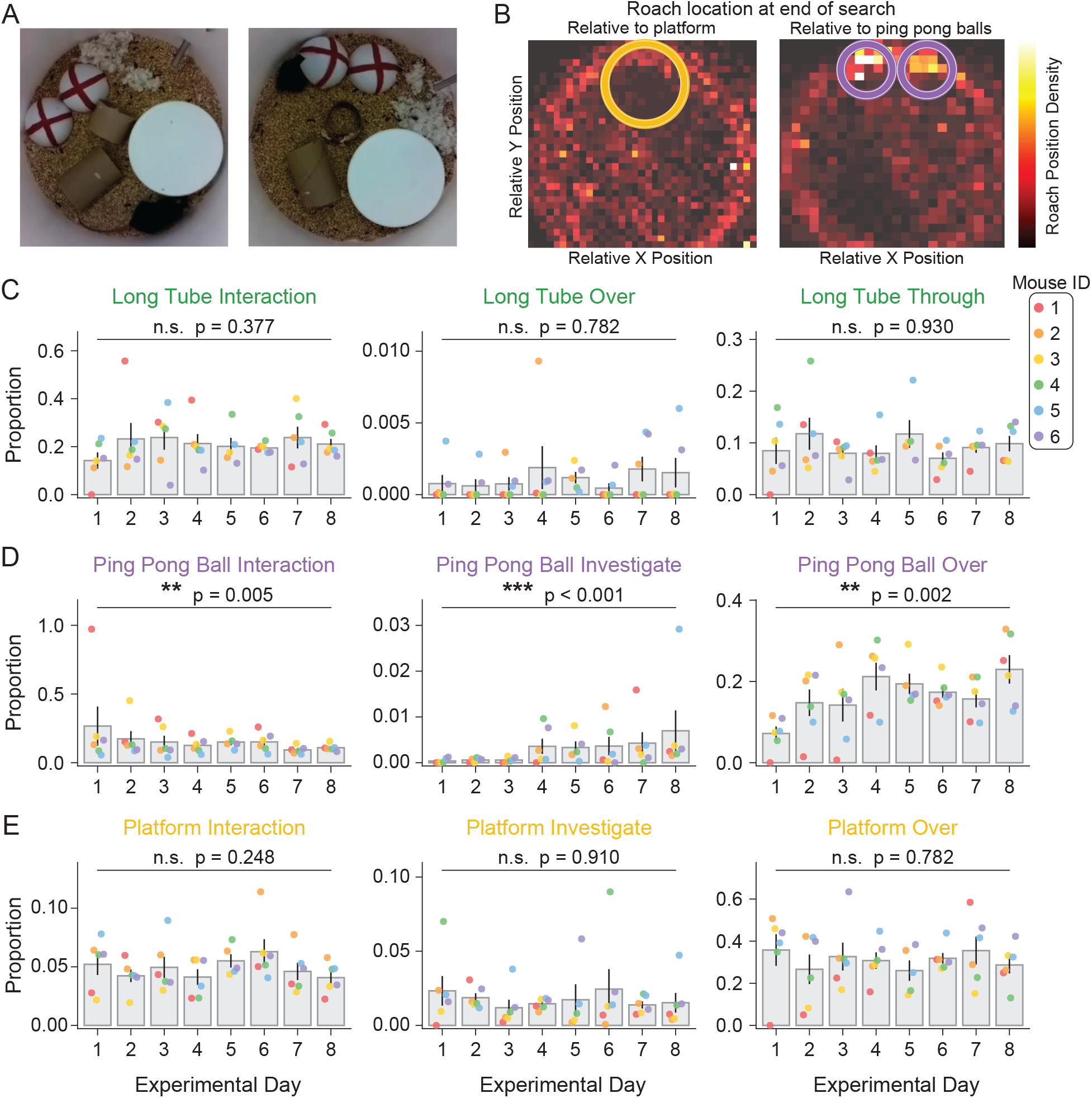
During search, mice learn to preferentially engage objects likely to uncover prey. **A**. Example video frames of platform (left) and ping-pong ball (right) investigations during prey capture, illustrating how ping-pong-ball investigation more readily exposes a concealed roach and re-initiates pursuit. **B**. Heatmaps of roach position density at the end of mouse searches, shown relative to the platform (left) and the ping-pong balls (right). Roaches were rarely located near the platform but were frequently located near the ping-pong balls when searches were successfully terminated, indicating that ping-pong-ball interactions are more likely to uncover prey. **C**. Proportion of long-tube-directed frames assigned to each long tube behavior (interaction, over, through) during the search state, across experimental days. For each animal on each day, the proportion was computed as the number of frames classified as a given behavior divided by the total number of object-directed frames during search that day. Bar shows the animal-level mean of these proportions; scatter points show individual animals (Mouse ID legend, top right). Brackets indicate the day-1-versus-day-8 contrast from a binomial generalized linear mixed model (cbind(behavior, other) ~ day + (1|animal/day)), with *p*-values FDR-corrected across the nine behaviors (n.s., not significant; ^∗∗^*p* < 0.01, ^∗∗∗^*p* < 0.001). **D**. Same as C, but for ping-pong ball behaviors (interaction, investigate, over). Over the course of learning, ping-pong-ball engagement shifted significantly: mice significantly decreased the proportion of interactions but increased investigations and over actions. **E**. Same as C, but for platform behaviors (interaction, investigate, over).

## DISCUSSION

Animals must adapt their behavior to environments in which resources are sparse, hidden, and structured by physical constraints. For purposes of experimental tractability, these conditions are often deliberately removed from the controlled paradigms typically used to study prey capture as an ethologically relevant behavior. To recover *ecologically* relevant complexity as a necessary component of ethologically relevant cognition (while retaining experimental tractability), we developed an enriched prey capture task, reproducible with inexpensive materials, alongside a framework for analyzing behavior in explicit relation to environmental structure. Using these tools, we show that mice learn effectively in cluttered environments not by refining pursuit itself, but by reorganizing how they search. Across analyses of trajectories, kinematics, and object-referenced behaviors, learning is expressed primarily as changes in search strategy, while pursuit remains comparatively stable. It is important to note that the mice studied here were already expert hunters, having reached asymptotic performance in the sparse arena weeks before objects were introduced. Despite this prior consolidation, the addition of environmental complexity revealed an entire stratum of memory-dependent, value-guided search strategy that sparse-arena performance neither required nor exposed. This dissociation carries a general implication: an artificially sparse environment can support expert performance while failing to engage the cognitive machinery that defines the same categorical behavior in nature. Performance in such reduced paradigms therefore cannot be assumed to index the strategic and mnemonic processes that the behavior recruits in its evolutionary context.

The sensorimotor control of pursuit and its concomitant motivation, which much of the prey capture literature has emphasized^16,17,40,41^, are undoubtedly crucial for hunting and capturing prey; our results indicate, however, that it is not where the signatures of learning in complex environments principally appear. The reorganization of search that we observe instead comprises a coherent set of changes in how mice explore and exploit the environment: increased movement variability, reduced fragmentation of behavioral states, more structured and less repetitive trajectories, and a growing preference for the object interactions most likely to reveal prey. These adaptations occur predominantly during search rather than pursuit, and together they reflect the progressive refinement of strategies for locating hidden prey under uncertainty.

A defining feature of this task is that environmental structure directly shapes behavior. Objects constrain movement, modulate visibility, and provide opportunities for concealment, thereby coupling predator and prey dynamics. Rather than treating the environment as uniform, mice exhibit object-specific patterns of behavior that reflect the functional properties of individual features (such as whether an object can be displaced to flush prey). By explicitly quantifying object-referenced actions, our framework links behavioral structure to environmental context and task demands, providing a means to analyze how behavior is organized in relation to environmental structure.

Our modeling results show that a small set of mechanisms can account for many of these observations. A minimal agent-based framework incorporating visibility-dependent pursuit, prey hiding, short-term spatial memory, and object-specific valuation was sufficient to reproduce key features of behavior in the enriched environment. Short-term memory reduces redundant revisitation and generates structured search sequences, while object-specific valuation aligns search effort with the distribution of prey. These results show that complex behavioral organization can emerge from a small set of interpretable constraints, without detailed motor control or high-dimensional policy representations.

Methodologically, this work pairs a compact, interpretable ethogram with hierarchical classification based on pose and object features, allowing actions to be analyzed explicitly in relation to environmental structure and task demands. Unsupervised approaches excel at discovering recurring patterns in high-dimensional behavior, but because they are not referenced to the environment, behavior-environment coupling remains implicit. By tying each action to its environmental context and consequences, our framework makes that coupling explicit, so that behavioral structure can be interpreted in terms of its role in the task. Applying it revealed that learning in this task is selective rather than uniform: instead of broadly increasing exploration, mice came to favor the specific interactions that expose hidden prey. Learning therefore operates not only over where animals search but over how they engage particular features of the environment to extract information—an adaptation to the causal structure of the task.

Several limitations should be noted. The prey (red runner) is not merely a stimulus but a co-agent whose hiding behavior is itself part of what the mouse learns; because we observed the roach’s position only intermittently and could not control its choices, our estimate of the hiding distribution the mouse adapts to is necessarily approximate, which bounds how precisely the value-matching account can be tested. Relatedly, the object set used here samples only a small portion of natural environmental complexity; our framework predicts that different configurations should elicit different search strategies, a prediction that future work can test directly by manipulating object layout and affordances. Two further caveats are more technical: some labeling ambiguity is unavoidable for behaviors that transition on sub-second timescales, and the agent-based model is intentionally minimal, capturing the logic of visibility, hiding, memory, and value rather than the full richness of sensory processing or motor control. None of these considerations alter the central finding that learning in this task is expressed primarily through the reorganization of search.

More broadly, our findings highlight the importance of memory and experience in shaping adaptive behavior. The convergence of search behavior toward the prey’s hiding distribution implies the formation of object-specific expectations, extending beyond short-term avoidance of recently visited locations. These results suggest that successful performance depends on integrating information across timescales to guide future exploration, consistent with the engagement of longer-term memory and value-based decision processes.

This emphasis distinguishes the present paradigm from traditional prey capture assays, which are often dominated by the rapid, sensorimotor aspects of pursuit. By requiring animals to infer likely prey locations and adapt their behavior across trials, this task places greater demands on memory, valuation, and strategy formation, and it reads out those processes directly through the organization of search. As such, it offers a behavioral window onto cognitive functions that low-dimensional assays—open-field, spatial mazes, and similar tests—are poorly positioned to resolve, particularly the strategic and mnemonic processes that may be altered well before they produce gross deficits in conventional measures. This makes it a candidate tool across a range of disease, aging, and neurodevelopmental models in which higher-order cognition is affected. Deficits in memory-guided search, in the allocation of exploration, or in learning object-specific value could be especially informative in disorders such as Alzheimer’s disease, where cognitive and decision-making processes are disrupted.

In summary, adaptive behavior in complex environments emerges through the progressive organization of search under environmental constraints. Rather than refining isolated actions, animals learn to structure their interaction with the environment in ways that efficiently reveal hidden resources. This perspective reframes behavior as an emergent property of agent–environment interactions and provides a foundation for studying learning in naturalistic contexts.

**Figure S1:**
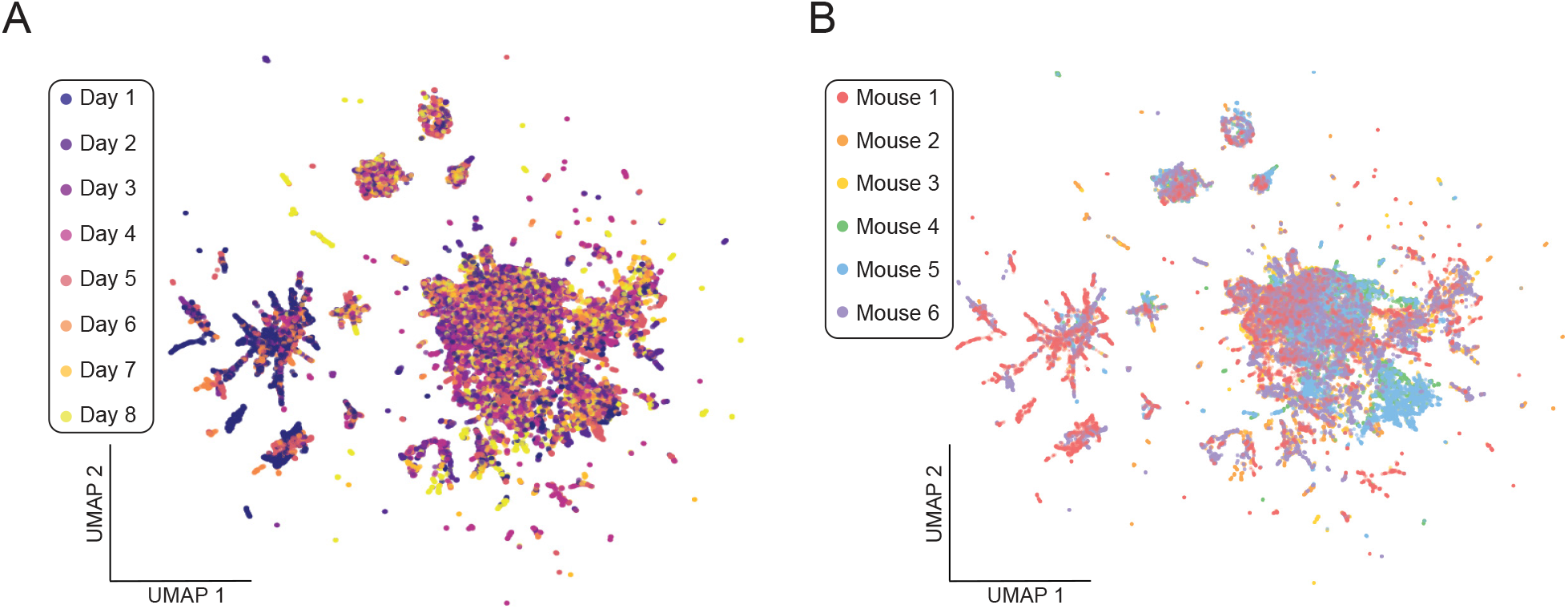
UMAP visualization of movement features. **A**. UMAP projection of mouse trajectories through the cluttered arena, colored by experimental day. The embedding revealed two prominent clusters, one enriched for early trials and another enriched for later trials. **B**. Same as in A, but colored by individual animal. The two clusters were differentially occupied by individual animals. Note that UMAP coordinates do not preserve distances, so the embedding is shown for visualization only; quantification of learning-related change is provided by the principal component analysis in Fig. 1G-I.

**Figure S2:**
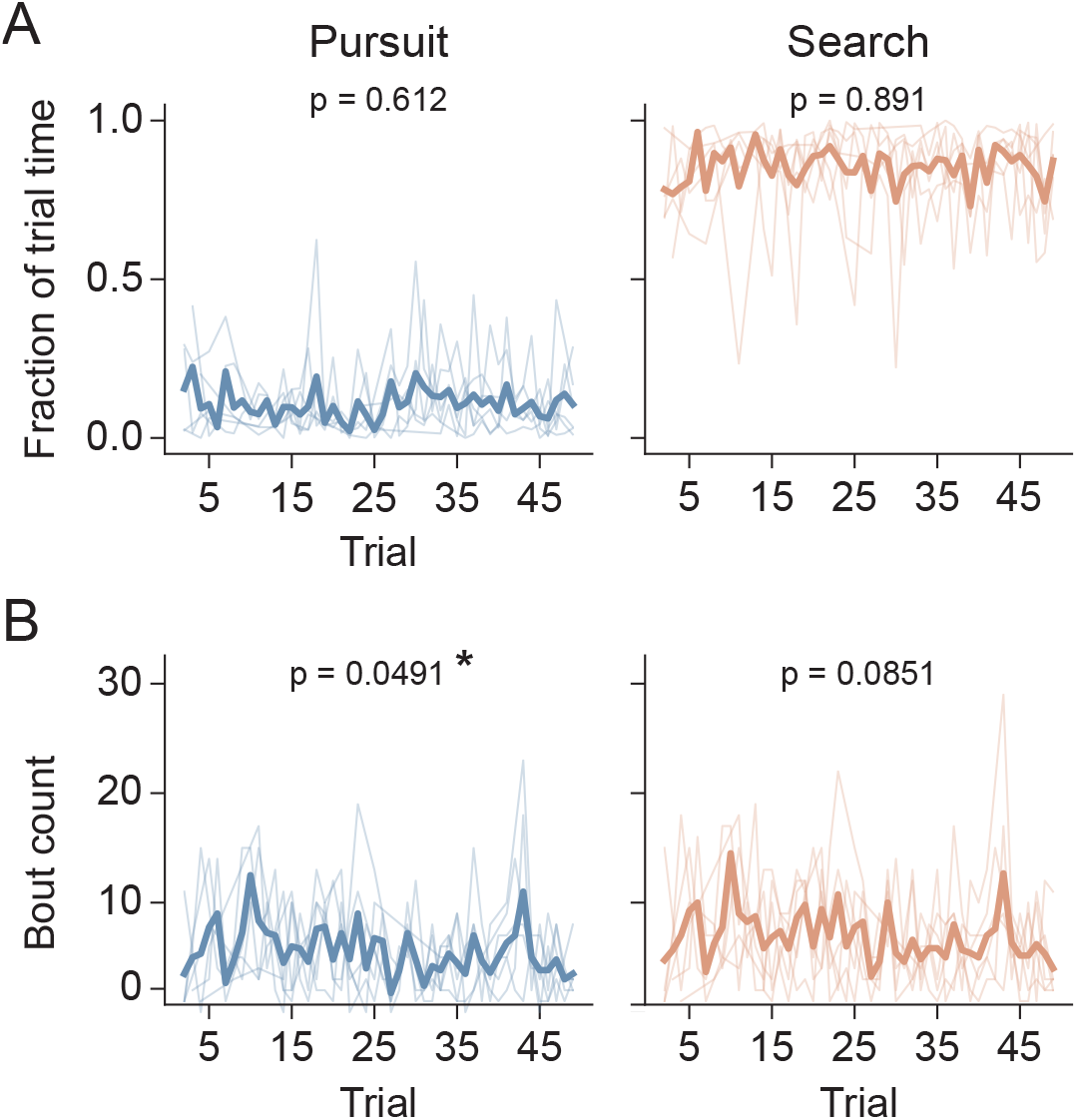
Pursuit and search bout structure is largely stable across trials. Bout statistics for pursuit (left, blue) and search (right, orange) states over the course of all trials. Thin lines show individual animals; thick lines show the group mean. P-values are from linear mixed-effects models (feature ~ trial + (1|animal), with animal as a random effect), testing for a change across trials. **A**. Fraction of trial time spent in each state. Mice spent the majority of each trial in the search state and a small fraction in pursuit, with neither changing significantly across trials (pursuit: p = 0.612; search: p = 0.891). **B**. Bout count. The number of bouts per trial showed a marginal decrease across trials in the pursuit state (* p = 0.049) and a non-significant trend in the search state (p = 0.085).

**Figure S3:**
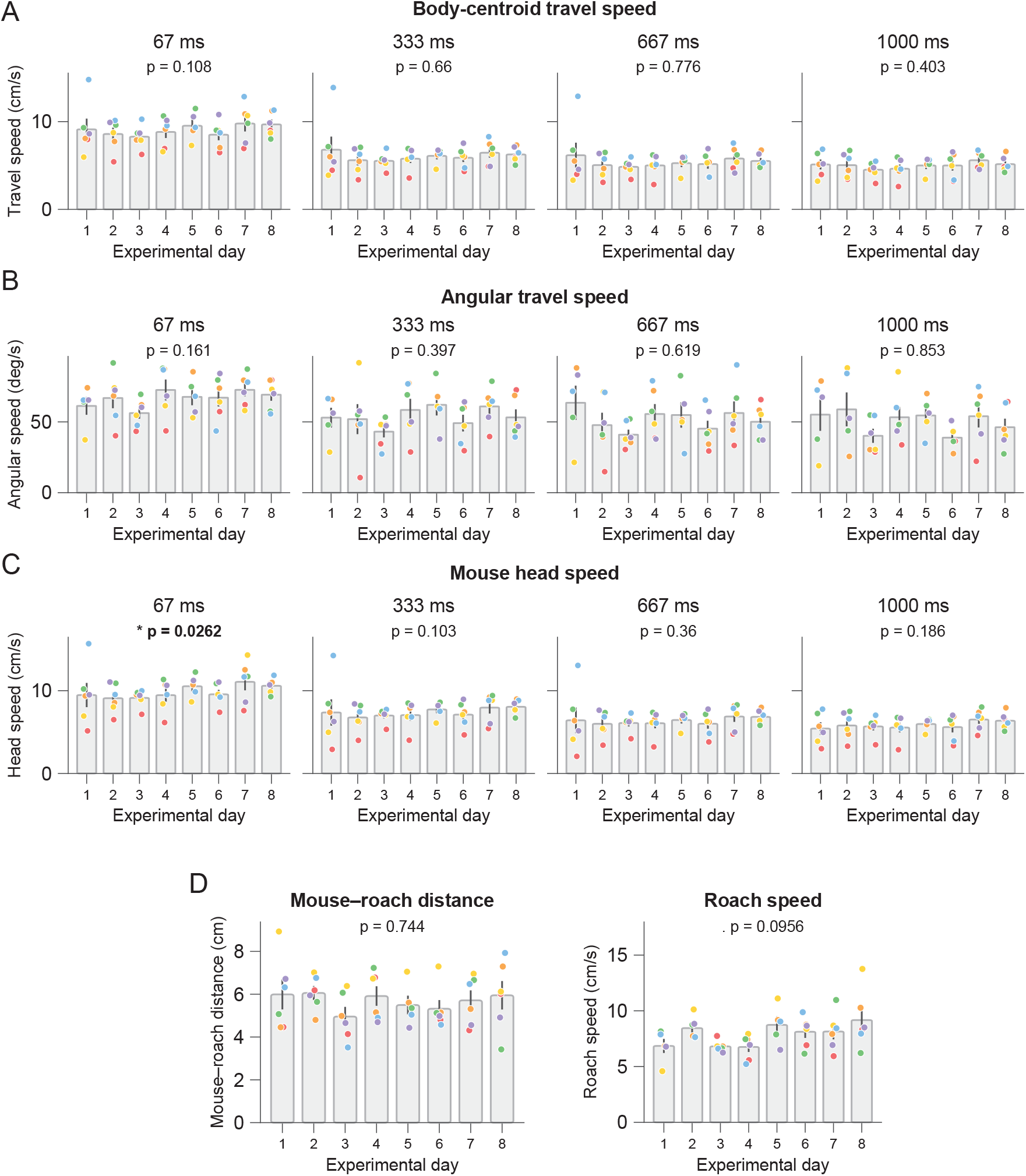
Movement kinematics during pursuit are largely stable across days of enriched prey capture. Each panel shows a kinematic feature averaged per animal on each experimental day, computed over four temporal windows (67, 333, 667, and 1000 ms; corresponding to 1, 5, 10, and 15 frames at 15 fps). Each dot represents the per-animal mean; bars show the group mean *±* SEM. Per-animal coloring is consistent across panels. P-values are from linear mixed-effects models (feature experimental ~ day + (1|animal), with animal as a random effect), comparing Day 1 vs. Day 8. **A**. Body-centroid travel speed. Translational speed of the body centroid did not change significantly across days at any timescale (all p *>* 0.1). **B**. Angular travel speed. The rate of change of heading direction did not differ significantly across days at any timescale (all p *>* 0.15). **C**. Mouse head speed. Head speed increased modestly across days at the shortest (67 ms) timescale (* p = 0.026) but not at longer timescales (p *>* 0.1). **D**. Mouse–roach distance (left) and roach speed (right). Neither the mean distance between mouse and roach (p = 0.744) nor roach travel speed (p = 0.096) changed significantly across days.

**Figure S4:**
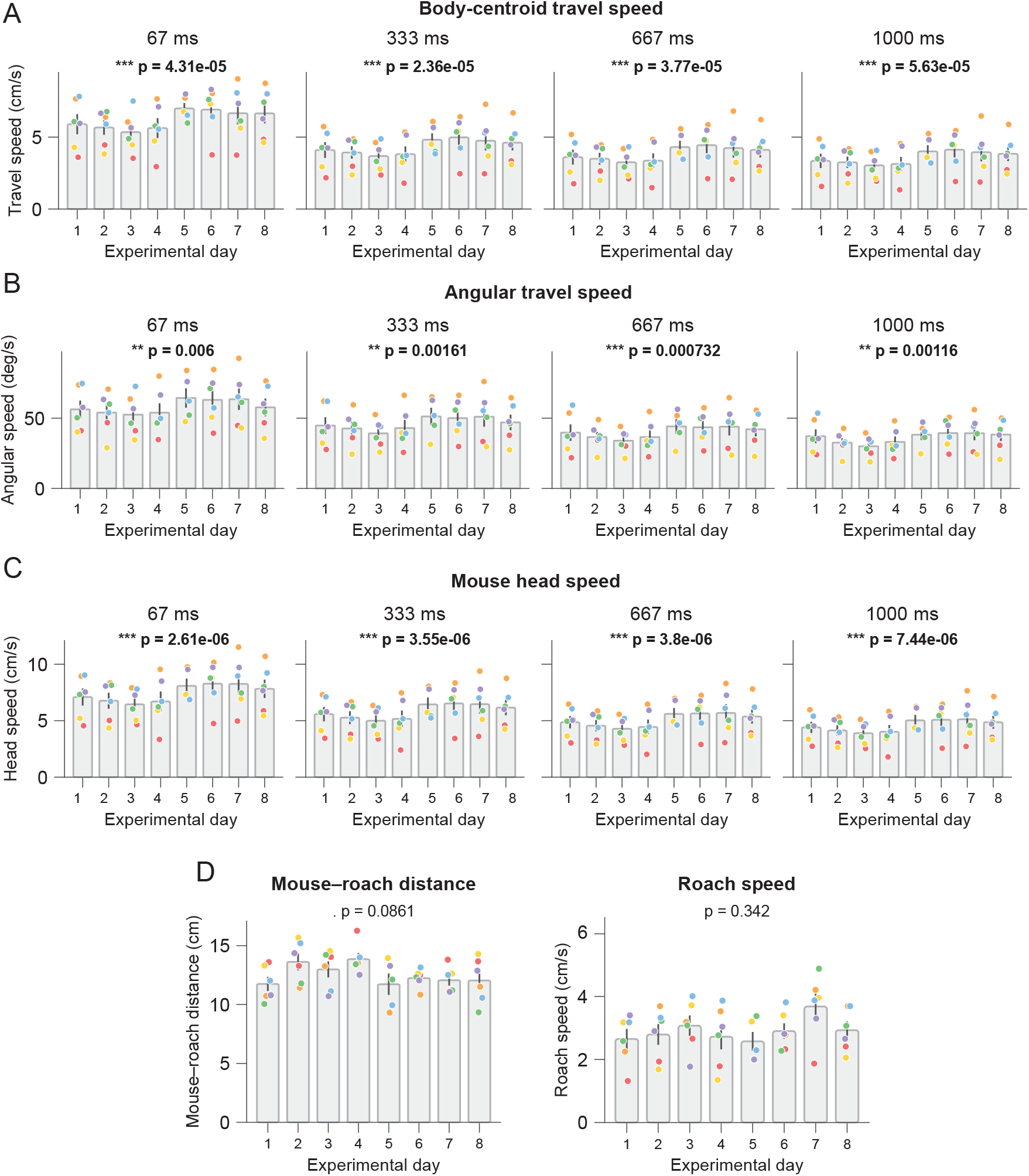
Movement kinematics during search change significantly across days of enriched prey capture. Each panel shows a kinematic feature averaged per animal on each experimental day, computed over four temporal windows (67, 333, 667, and 1000 ms; corresponding to 1, 5, 10, and 15 frames at 15 fps). Each dot represents the per-animal mean; bars show the group mean *±* SEM. Per-animal coloring is consistent across panels. P-values are from linear mixed-effects models (feature experimental ~ day + (1|animal), with animal as a random effect), comparing Day 1 vs. Day 8. **A**. Body-centroid travel speed increased significantly across days at all timescales (all p < × 10^−4^; 67 ms: p = 4.31 × 10^−5^; 333 ms: p = 2.36 × 10^−5^; 667 ms: p = 3.77 × 10^−5^; 1000 ms: p = 5.63 × 10^−5^). **B**. Angular travel speed changed significantly across days at all timescales (67 ms: p = 0.006; 333 ms: p = 0.00161; 667 ms: p = 0.000732; 1000 ms: p = 0.00116). **C**. Mouse head speed increased significantly across days at all timescales (all p < × 10^−5^; 67 ms: p = 2.61 × 10^−6^; 333 ms: p = 3.55 × 10^−6^; 667 ms: p = 3.8 × 10^−6^; 1000 ms: p = 7.44 × 10^−6^). **D**. Neither mouse–roach distance (left, p = 0.086) nor roach speed (right, p = 0.342) changed significantly over the course of experimental days.

**Figure S5:**
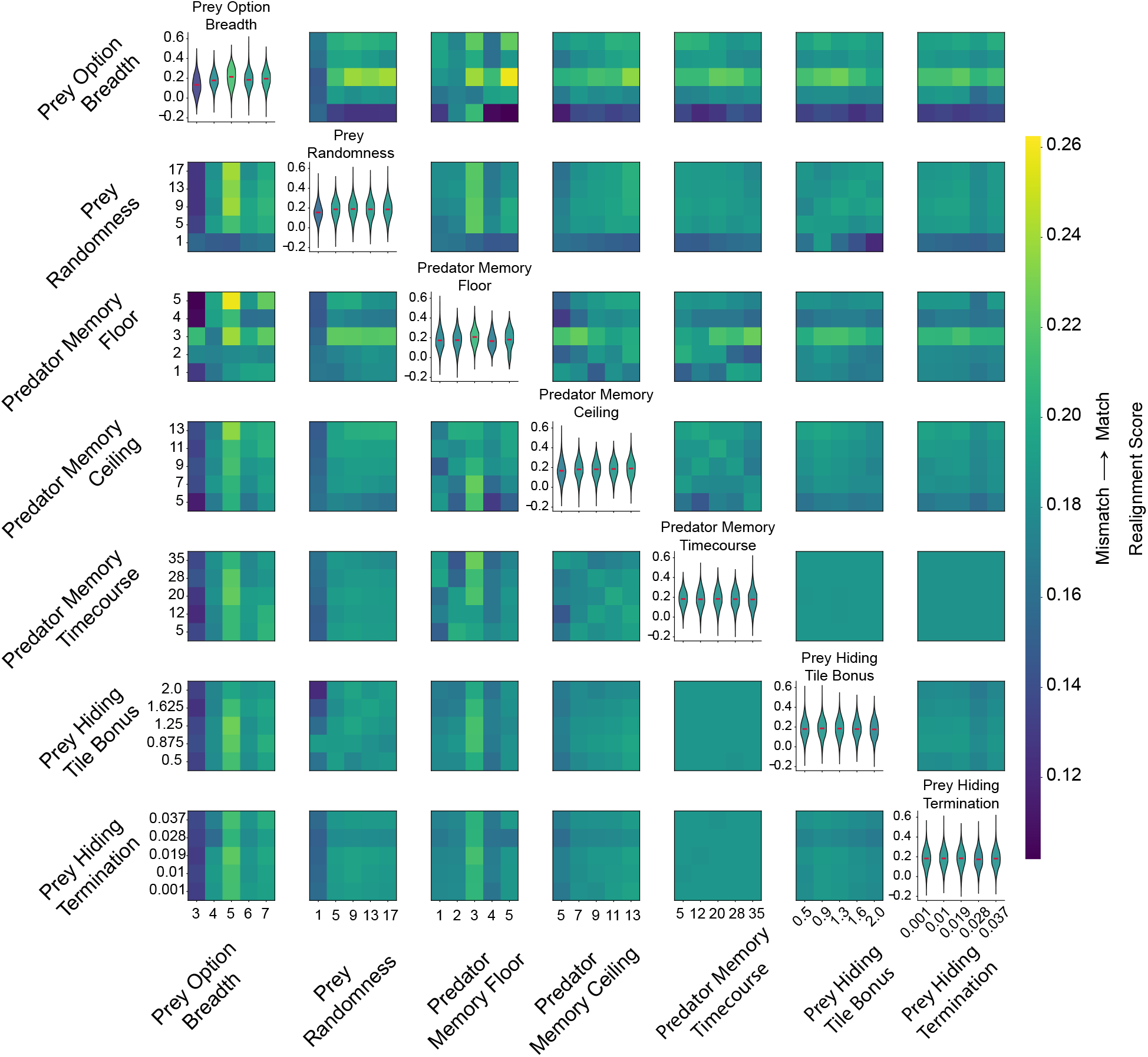
Predator search-memory and prey-decision hyperparameters jointly govern match/mismatch tile-occupancy realignment. Pairwise summary of a 5^7^ = 78,125-combination grid sweep over seven hyperparameters governing the prey decision policy and predator search memory: `prey_top_n` (number of top-scoring candidate flee moves the prey’s tree-search planner samples from), `prey_temperature` (softmax temperature controlling how uniformly the prey samples among those candidates), `sterm_mem_min` (predator search-memory cooldown floor, in turns, immediately after a sighting), `sterm_mem_max` (asymptotic ceiling the cooldown approaches during a prolonged search), `sterm_mem_tau` (timescale, in turns, over which the cooldown grows from floor to ceiling), `prey_hide_bonus` (bonus added to a candidate flee move’s score for leading toward a concealment-capable tile), and `pexpire` (per-turn probability a hidden prey spontaneously becomes visible again). Off-diagonal cells show the color-mapped Mismatch → Match Realignment Score – the divergence between roach and mouse object-tile occupancy proportions during search under a deliberately uninformed (*mismatch*) predator search prior, minus the same divergence under the correct (*match*) prior – averaged over all grid combinations sharing that pair of row/column parameter values (marginalizing over the other five parameters); higher values indicate a larger improvement in roach/mouse tile-occupancy alignment when the predator’s search prior is correctly informed. Diagonal panels show the full marginal distribution (violin) of the same score across all combinations for each tested value of that parameter alone, with the per-value mean (red tick) colored by its position on the shared colorbar. Colorbar limits are clipped to the [20th, 80th] percentile of all cell values across the sweep to enhance contrast among the bulk of the grid.

**Figure S6:**
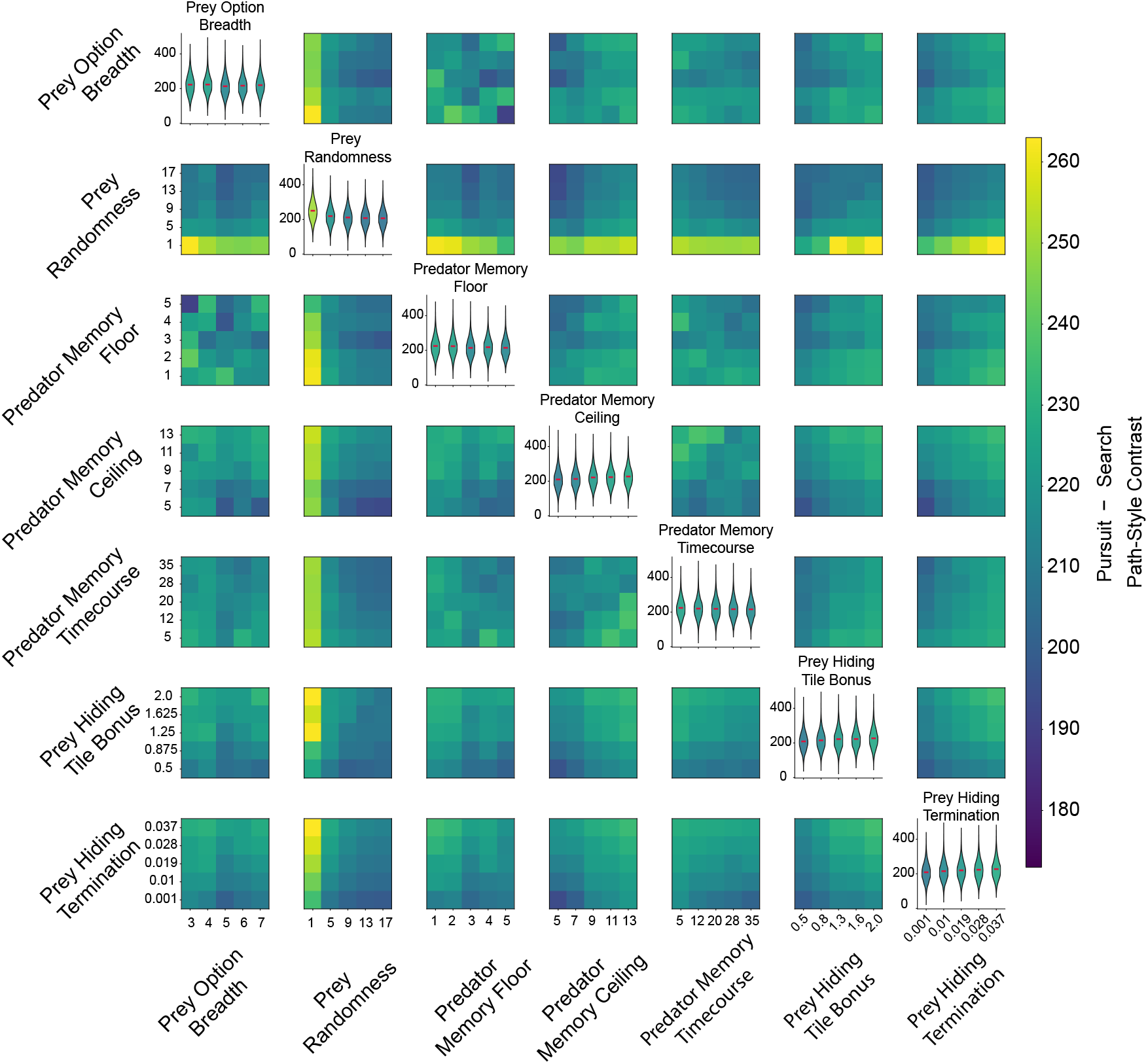
The same hyperparameters shape a Search−Pursuit path-style contrast. Pairwise summary of the same 5^7^ = 78,125-combination grid sweep over `prey_top_n`, `prey_temperature`, `sterm mem min`, `sterm_mem_max`, `sterm_mem_tau`, `prey_hide_bonus`, and `pexpire` (as defined in Fig. S5), restricted to the ecologically valid *match* condition. Off-diagonal cells show the color-mapped Search−Pursuit Path-Style Contrast – `search_net_paths` (loop count minus back-and-forth count during the predator’s search state) minus `pursuit_net_paths` (the same quantity during pursuit) – averaged over all grid combinations sharing that pair of row/column parameter values (marginalizing over the other five parameters); higher values indicate a combined preference for looping trajectories during search together with direct back-and-forth trajectories during pursuit, the qualitatively expected behavioral split between the two states. Diagonal panels show the full marginal distribution (violin) of the same contrast across all combinations for each tested value of that parameter alone, with the per-value mean (red tick) colored by its position on the shared colorbar. Colorbar limits are clipped to the [20th, 80th] percentile of all cell values across the sweep to enhance contrast among the bulk of the grid.

**Figure S7:**
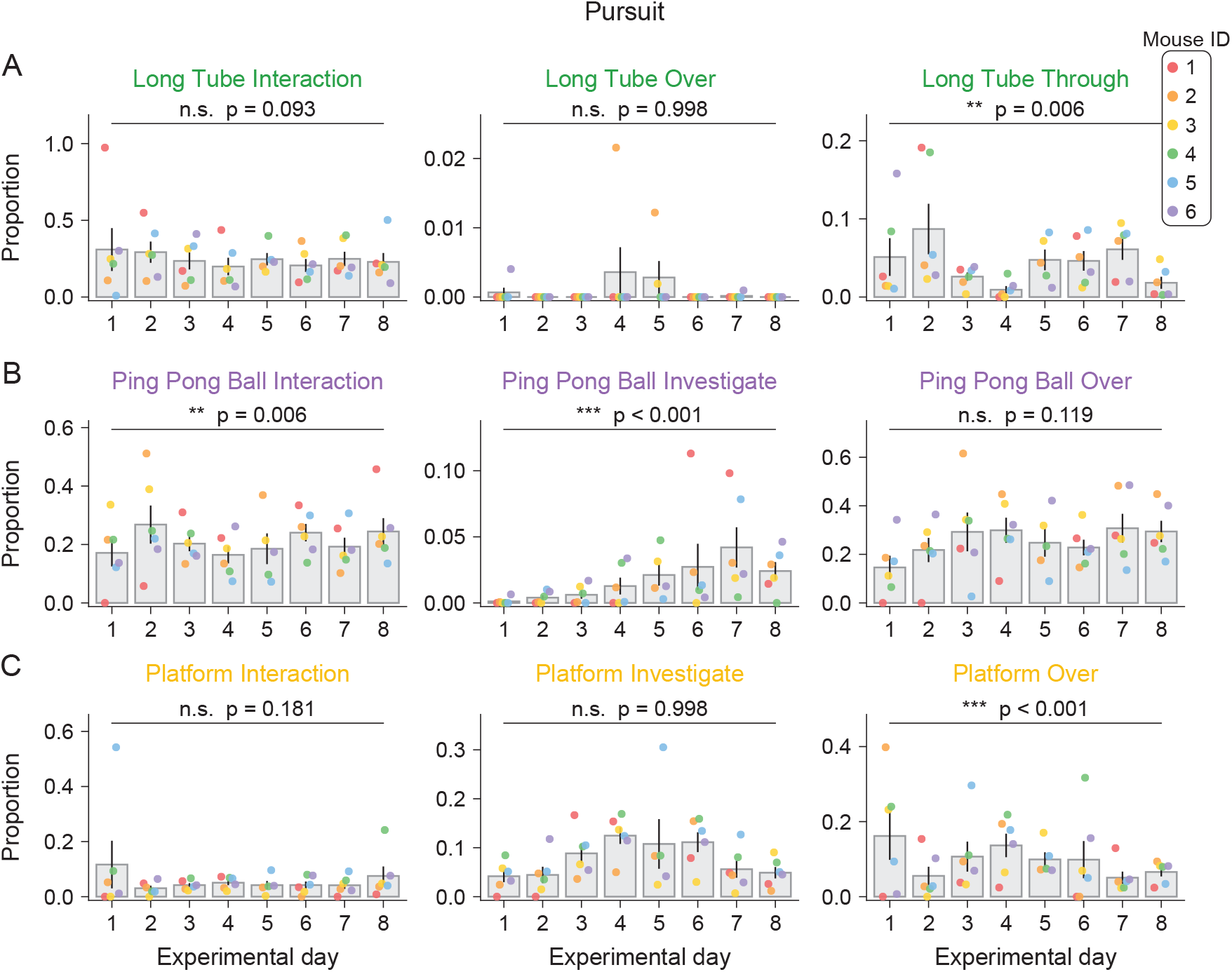
Object-directed behavior during pursuit shifts across learning. **A**. Proportion of long-tube-directed frames assigned to each long tube behavior (interaction, over, through) during the pursuit state, across experimental days. For each animal on each day, the proportion was computed as the number of frames classified as a given behavior divided by the total number of object-directed frames during pursuit that day. Bars show the animal-level means of these proportions; scatter points show individual animals (Mouse ID legend, top right). Brackets indicate the day-1-versus-day-8 contrast from a binomial generalized linear mixed model (cbind(behavior, other) ~ day + (1|animal/day)), with *p*-values FDR-corrected across the nine behaviors (n.s., not significant; ^∗∗^*p* < 0.01, ^∗∗∗^*p* < 0.001). Of the long tube behaviors, only long-tube-through changed significantly across learning (** p = 0.006), while interaction (p = 0.093) and over (p = 0.998) did not. **B**. Same as A, but for ping-pong ball behaviors (interaction, investigate, over). Ping-pong-ball engagement shifted significantly over learning: mice changed the proportion of interactions (** p = 0.006) and investigations (*** p < 0.001), while over actions did not change significantly (p = 0.119). **C**. Same as A, but for platform behaviors (interaction, investigate, over). Platform-over actions changed significantly across learning (*** p < 0.001), while interaction (p = 0.181) and investigate (p = 0.998) did not.

**Table 1:** Hyperparameters used for each classification model.

| Model | Hyperparameters |
| --- | --- |
| Hierarchical XGBoost | **General behavior:** learning_rate=0.0599585914; max_depth=8; max_leaves=46; min_child_weight=1.7996666326; subsample=0.5337114619; colsample_bytree=0.8419617420; gamma=2.8624687049; n_estimators=520.<br>**PVC/cap behavior:** learning_rate=0.1257324308; max_depth=5; max_leaves=29; min_child_weight=7.4205703222; subsample=0.9990035994; colsample_bytree=0.7827709541; gamma=2.9476603104; n_estimators=590.<br>**PPBall behavior:** learning_rate=0.2294966354; max_depth=8; max_leaves=25; min_child_weight=9.0849077244; subsample=0.7329747878; colsample_bytree=0.5151673153; gamma=2.3420394159; n_estimators=1000.<br>**LTube behavior:** learning_rate=0.0972519032; max_depth=7; max_leaves=46; min_child_weight=1.3043964612; subsample=0.6677810473; colsample_bytree=0.9707083872; gamma=3.6287946899; n_estimators=670.<br>For all component classifiers: random_state=42. |
| Flat XGBoost | learning_rate=0.0599585914; max_depth=8; max_leaves=46; min_child_weight=1.7996666326; subsample=0.5337114619; colsample_bytree=0.8419617420; gamma=2.8624687049; n_estimators=520; random_state=42. |
| Hierarchical logistic regression | **General behavior:** C=0.2029426772; penalty=l2; solver=saga.<br>**PVC/cap behavior:** C=6.0864091301; penalty=l1; solver=saga.<br>**PPBall behavior:** C=9.9699022269; penalty=l2; solver=saga.<br>**LTube behavior:** C=7.4395288849; penalty=l1; solver=saga.<br>For all component classifiers: max_iter=1000; n_jobs=-1; random_state=42. |
| Flat logistic regression | C=2.5177425639; penalty=l2; solver=saga; max_iter=1000; n_jobs=-1; random_state=42. |
| Hierarchical random forest | **General behavior:** n_estimators=420; max_depth=20; min_samples_split=7; min_samples_leaf=2; max_features=sqrt; criterion=entropy; bootstrap=true.<br>**PVC/cap behavior:** n_estimators=320; max_depth=20; min_samples_split=14; min_samples_leaf=6; max_features=sqrt; criterion=gini; bootstrap=true.<br>**PPBall behavior:** n_estimators=180; max_depth=20; min_samples_split=11; min_samples_leaf=3; max_features=sqrt; criterion=entropy; bootstrap=true.<br>**LTube behavior:** n_estimators=50; max_depth=15; min_samples_split=2; min_samples_leaf=4; max_features=log2; criterion=gini; bootstrap=true.<br>For all component classifiers: n_jobs=-1; random_state=42. |
| Flat random forest | n_estimators=310; max_depth=19; min_samples_split=3; min_samples_leaf=3; max_features=sqrt; criterion=entropy; bootstrap=true; n_jobs=-1; random_state=42. |

Table 1 – continued from previous page
| Model | Hyperparameters |
| --- | --- |
| Hierarchical multilayer perceptron | The same settings were used for each component classifier: `hidden_layer_sizes=(64,); activation=relu; solver=adam; alpha=0.0001; batch_size=auto; learning_rate=constant; learning_rate_init=0.001; max_iter=1000; early_stopping=false; random_state=42.` |
| Flat multilayer perceptron | `hidden_layer_sizes=(64,); activation=relu; solver=adam; alpha=0.0001; batch_size=auto; learning_rate=constant; learning_rate_init=0.001; max_iter=1000; early_stopping=false; random_state=42.` |
| Hierarchical mixture of experts | The same settings were used for each component classifier: `num_experts=1.` Each expert and gating network was an MLP with `hidden_layer_sizes=(64,); activation=relu; solver=adam; learning_rate_init=0.001; max_iter=1000; random_state=42.` |
| Flat mixture of experts | `num_experts=1.` Each expert and the gating network was an MLP with `hidden_layer_sizes=(64,); activation=relu; solver=adam; learning_rate_init=0.001; max_iter=1000; random_state=42.` |
| Hierarchical Transformer, 512-unit feedforward layer | The same settings were used for each component classifier: `d_model=128; nhead=4; num_encoder_layers=2; num_decoder_layers=2; dim_feedforward=512; dropout=0; batch_first=true; sinusoidal positional encoding; feature rotation enabled; epochs=100; learning_rate=0.001; Adam optimizer; cross-entropy loss; full-batch training; random_state=42.` |
| Flat Transformer, 512-unit feedforward layer | `d_model=128; nhead=4; num_encoder_layers=2; num_decoder_layers=2; dim_feedforward=512; dropout=0; batch_first=true; sinusoidal positional encoding; feature rotation enabled; epochs=100; learning_rate=0.001; Adam optimizer; cross-entropy loss; full-batch training; random_state=42.` |
| Hierarchical Transformer, 256-unit feedforward layer | The same settings were used for each component classifier: `d_model=128; nhead=4; num_encoder_layers=2; num_decoder_layers=2; dim_feedforward=256; dropout=0; batch_first=true; sinusoidal positional encoding; feature rotation enabled; epochs=100; learning_rate=0.001; Adam optimizer; cross-entropy loss; full-batch training; random_state=42.` |
| Flat Transformer, 256-unit feedforward layer | `d_model=128; nhead=4; num_encoder_layers=2; num_decoder_layers=2; dim_feedforward=256; dropout=0; batch_first=true; sinusoidal positional encoding; feature rotation enabled; epochs=100; learning_rate=0.001; Adam optimizer; cross-entropy loss; full-batch training; random_state=42.` |

## METHODS

### Animal Husbandry

We used six (3 ♂, 3 ♀) C57BL/6 mice (Mus musculus) for all experiments described throughout. All mice were 3.9 months old at the time of starting the first set of experiments. Prior to prey capture experiments, mice were housed in an enriched environment and kept on a 12:12 h light:dark cycle with ad libitum access to food and water. We pooled data between male and female mice, as we observed no differences in predation between the sexes.

For prey, we used adult Turkestan red runner cockroaches (Supplier: Caribbean Mealworms) weighing 0.05 to 0.20 g. Roaches were randomly selected then weighed before being used for trials.

All procedures were performed in accordance with protocols approved by the Washington University in Saint Louis Institutional Animal Care and Use Committee, following guidelines described in the US National Institutes of Health Guide for the Care and Use of Laboratory Animals.

### Prey Capture Assay

During initial prey capture training in an unenriched arena, mice were trained individually to hunt red runner cockroaches (Shelfordella lateralis) in circular home-cage chambers (36.83 cm H × 25.87 cm D). During prey capture testing, chambers were empty aside from bedding.

Before the first day of trials, mice were acclimated to cockroaches by placing three insects in the chamber for two consecutive nights during the 12 h dark period. On the night before the first day of testing, mice were food deprived overnight for 15 h. On the first day of trials, at 9:30 AM (ZT 2.5), six cockroaches were introduced sequentially, with each new cockroach added after the preceding trial ended either by capture or by timeout, defined as *>*30 min without capture. Following these six trials, mice were provided ad libitum access to food and water until being food deprived again in advance for the following day of testing. Although capture times reached an asymptotic, expert-level performance within the first four days, training continued for 10 weeks to produce expert hunters under these conditions. On the day before enriched prey capture testing, trials were again conducted in the empty arena to confirm mice maintained expert-level performance.

During enriched prey capture testing, mice underwent an eight-day protocol. These enriched arenas contained a fixed set of repurposed objects: 1) a large PVC pipe cap, used as a platform; 2) a bathroom tissue roll core, used as a long tube; 3) a short segment of a bathroom tissue roll core, used as a short tube; and 4) two halves of a jumbo ping-pong ball. These objects were removed from the chamber during times outside of prey capture trials and were replaced with food pellets and nestlets.

### Video recording and Animal/Object tracking

Video recordings were performed using e3Vision cameras and the Watch Tower (White Matter LLC) system. A camera was placed directly above the center of each arena. Videos were recorded at 15 frames per second.

To quantify the position and posture of mice, we used DeepLabCut^36^, a convolutional network for markerless pose estimation. We tracked six keypoints on each mouse: the snout, left ear, right ear, shoulder, mid-back (spine), and base of the tail. A model previously trained on mice in the unenriched arena was used to estimate keypoint positions. Occlusions introduced by objects in the enriched arena (primarily when mice passed through the long tube) were infrequent; undetected keypoints on a given frame were treated as missing and took on a default value (which was excluded from statistical analyses but included for classification with machine-learning models). For analyses requiring a single mouse position, we used a “mean head” point defined as the average of the snout, left ear, right ear, and shoulder positions, ignoring missing keypoints.

The roach and arena objects were localized using convolutional object-detection models based on the YOLO architecture^37^. One model tracked the roach as a single point at the center of its back; a second model continuously localized the arena objects—the platform, long tube, short tube, two ping-pong ball halves, and water spout—each within a bounding box, with object position taken as the bounding-box center. Because the roach was frequently occluded by objects owing to its small size, often remaining hidden for tens of seconds, we supplemented model predictions with manual annotation in which a scorer labeled the roach’s position on one of every 15 frames (one labeled frame per second) to constrain and correct model output. When the roach was truly occluded, its last observed position was assumed to persist.

### Behavior Classification

All trials were segmented into three behavioral states—pursuit, search, and abandonment—by a panel of three independent human scorers. Scorers achieved *>*90% pairwise agreement across all trials, and consensus labels were used for all subsequent state-based analyses.

To classify object-directed behaviors at frame resolution, we trained supervised classifiers on features derived from DeepLabCut pose estimates and YOLO object localizations: pairwise distances between mouse keypoints, distances from the snout and spine to each object, head- and body-orientation angles relative to each object, and mouse velocity at multiple temporal lags (1–45 frames). Features were min–max normalized to [0, 1] using training-set bounds, with missing values encoded as −2.

We defined a ten-category ethogram—interaction, investigate/through, and over behaviors for the platform, ping-pong balls, and long tube, plus a no-action category—and compared flat and hierarchical classification structures. The hierarchical model first predicted which object (if any) the mouse engaged, then predicted the behavior within that object group, combining stages by weighting each within-object probability by its object probability. We evaluated XGBoost, logistic regression, random forest, multilayer perceptron, mixture-of-experts, and transformer architectures, training on all but one held-out labeled video and quantifying performance as balanced accuracy. The hierarchical XGBoost model performed best and was used for all subsequent ethogram analyses.

### Kinematic analysis

Movement kinematics were computed at multiple temporal scales by measuring displacement over frame-lag windows of 1, 5, 10, and 15 frames (67, 333, 667, and 1000 ms at 15 fps) and dividing by the elapsed time. Body-centroid travel speed was computed from the mean position of the shoulder, spine, and tailbase keypoints; head speed from the head keypoints; and angular speed as the rate of change of heading direction. Speeds in Fig. 2A were normalized to the maximum value across the dataset. Pursuit and search states were compared using these metrics, and each metric was tested for changes across experimental days. Statistical comparisons used linear mixed-effects models with animal as a random effect (see Statistical Analyses).

### Dimensionality reduction and path analysis

To characterize the structure of mouse trajectories through the arena, we performed dimensionality reduction on path segments. Because aligning raw image (x, y) coordinates across animals and trials was unreliable—objects were subtly displaced by the mouse over the course of a trial, and object positions were only approximately reset between trials—we represented each position not in image coordinates but in a multidimensional coordinate system defined by the mouse’s distance to each object in the arena (platform, long tube, short tube, and ping-pong balls). This extends the intuition that, in an empty circular arena, a mouse’s position can be summarized by its distance to the wall, and preserves the spatial relationships between objects without requiring precise cross-trial registration.

Paths were analyzed over a range of timescales by extracting segments from 1 to 10 seconds in length. We computed pairwise distances between segments using dynamic time warping and projected them into low-dimensional spaces using multidimensional scaling. For each timescale, points corresponding to individual path segments were colored by continuous scalar descriptors (e.g., total distance traveled, mean and variance of speed) and by categorical descriptors (e.g., animal identity, experimental day) to identify the principal sources of variance in the low-dimensional embedding.

### Agent-based Modeling

#### Predator–prey model overview

We implemented the predator–prey model as a turn-based gridworld using the MiniGrid framework. The arena was a fixed 6 × 6 grid containing wall, open-floor, and object tiles. In the arena containing objects, four object types were positioned to approximate their relative locations in the experimental arena. A single predator and a single prey were initialized on randomly selected open tiles at the beginning of each episode. Each episode continued until the prey was captured or a maximum of 2,700 turns was reached.

At turn *t*, the predator and prey occupied positions 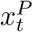 and 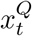, respectively. Both agents used egocentric action sets consisting of four cardinal moves, four diagonal moves, and a stationary action, denoted STAY. Illegal actions directed into walls produced no displacement. Distance was measured throughout as Chebyshev distance, *d*_∞_, consistent with the eight-connected adjacency of the grid. Each non-wall tile had a type

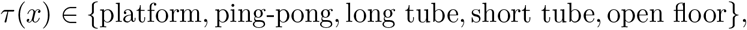

where open floor denoted an unoccupied tile and the remaining four types denoted the object categories, which differed in their concealment properties.

Both agents followed fixed heuristic policies and had no learning component. The predator’s behavioral state depended on whether the prey was visible. The prey was considered visible when it was unconcealed and within the predator’s egocentric field of view. When visible, the prey elicited pursuit; when it was not visible, the predator engaged in search.

#### Predator policy

When the prey was visible, the predator pursued it by selecting the legal action that minimized the resulting Chebyshev distance:

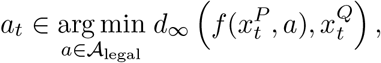

where *f* (*x, a*) denotes the position reached by taking action *a* from position *x*. Ties were resolved first in favor of non-diagonal moves and then at random. Any sighting of the prey reset the predator’s search memory to its baseline state.

When the prey was not visible, the predator selected among adjacent legal tiles according to a tile-type-specific search-value function, *V* (*τ*). Search values were transiently suppressed on recently visited tiles to produce novelty-seeking search. Each tile was assigned a cooldown, *C*_*t*_(*x*), when visited, and its effective value at turn *t* was

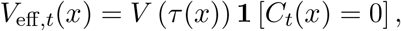

where **1**[*·*] denotes the indicator function. The predator selected

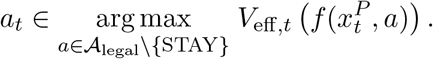

The STAY action was used only when movement in every direction was blocked.

The cooldown duration assigned to a visited tile increased as a function of the number of turns, *s*_*t*_, elapsed since the prey was last sighted:

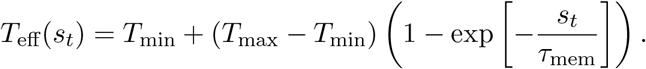

Active cooldowns decreased by one on each subsequent turn, and all cooldowns were reset to zero immediately when the prey was sighted. In all simulations reported here,

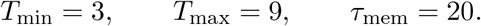

Thus, recently visited locations remained suppressed for progressively longer periods as the time since the previous prey sighting increased.

The tile-type value function implemented the experimental value-match manipulation. In the value-matched condition, the search value *V* of each object was set according to empirically derived per-object concealment probabilities, estimated from the observed distribution of prey hiding locations on the first day of enriched exposure:

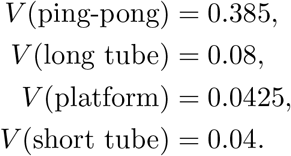

Open floor was assigned a small positive value, *V* (open floor) = 0.01, so that search was not completely restricted to object tiles.

In the value-mismatched condition, search values were uniform across object types:

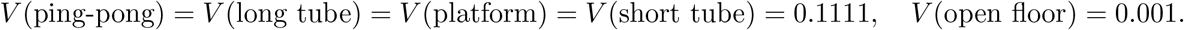

The value-matched and value-mismatched conditions differed only in these value assignments; all other aspects of the predator policy were identical.

#### Prey policy

While concealed, the prey followed a stationarity-biased random walk over actions that produced a legal displacement, together with the option to remain stationary:

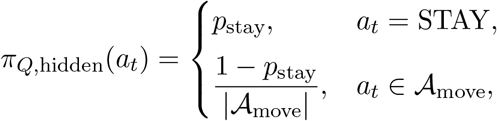

where *A*_move_ denotes the set of actions producing legal movement. We used *p*_stay_ ≈ 0.99, such that concealed prey were overwhelmingly immobile.

While visible, the prey evaluated its immediately available actions using a value function that combined evasion and concealment. The STAY action was excluded when the predator was within a Chebyshev distance of two tiles. Each candidate action *a* was scored as

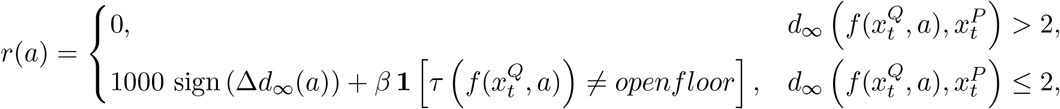

where

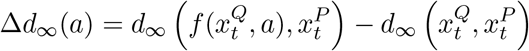

was the change in predator–prey distance produced by the candidate action. The concealment bonus was uniform across object types, with *β* = 0.5.

Up to the five highest-scoring legal actions were retained, and the executed action was sampled from a Boltzmann distribution over these candidates:

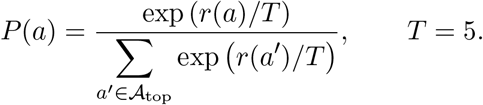

Because the evasion term was three orders of magnitude larger than the concealment term, the policy effectively selected an evasive action whenever one was available. Stochasticity primarily resolved ties among similarly evasive actions in favor of tiles offering concealment.

Independently of this action evaluation, whenever the predator was more than two tiles away, the selected action was replaced with STAY with probability 0.8. This override produced a strong immobility bias when the predator was not an immediate threat.

#### Concealment, detection, and capture

Concealment success depended on the tile occupied by the prey. When an unconcealed prey entered an object tile, it attempted to hide with the following object-dependent probabilities:

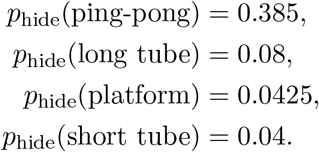

The prey could not conceal itself on open floor, so *p*_hide_(open floor) = 0.

A concealed prey could spontaneously become visible on any turn with probability

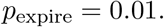

When the predator contacted a concealed prey, a search roll determined whether the prey was revealed. The search probability was

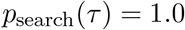

for all tile types in the present configuration. Contact with a concealed prey therefore always revealed it, and detection failure did not contribute stochasticity to the reported simulations.

Following contact with a revealed prey, whether the prey had been visible on arrival or had just been revealed through search, capture succeeded with a tile-dependent probability. Capture probability was 0.0588 on object tiles and 0.2 on open floor. This capture roll was the only remaining source of stochasticity between predator–prey contact and episode termination.

#### Arena and experimental conditions

We used two arena configurations that shared the same 6 × 6 wall boundary and model parameters but differed in their interior tile layouts and predator field-of-view sizes. The “with objects” condition contained the four object categories described above, positioned to approximate the relative arrangement of objects in the experimental arena, and used a 3 × 3 egocentric field of view.

The “without objects” condition replaced all interior object tiles with open floor and used a wider 5 × 5 field of view. Because *p*_hide_(*openfloor*) = 0, prey could not conceal themselves in this condition. Episodes therefore consisted of continuous pursuit interrupted only by transient loss of the prey from the predator’s field of view rather than by concealment.

Within the with-objects arena, we compared the value-matched and value-mismatched predator conditions to isolate the contribution of value-guided search (Fig. 5C). To match the size of the experimental cohort, we performed six independent simulation runs (one per random seed) per condition, treating run as the unit of replication in all analyses of simulated behavior, analogous to individual animals in the experimental data. All simulations were generated using the model’s internal predator and prey policies, without external control or policy learning.

#### Analysis of simulated behavior

Object-visit sequences generated during simulated search and pursuit were analyzed using the same procedures applied to the experimental behavioral data. Specifically, observed object-visit triplet frequencies and sample entropy were compared with null distributions generated from shuffled visit sequences (Fig. 4D,E). This analysis allowed the sequential structure of model behavior to be evaluated directly against that observed in mice.

#### Statistical Analyses

Unless otherwise noted, all statistical analyses used linear mixed-effects modeling using the lmer function from the lme4 package in R^42^, allowing us to test the potential influence of different fixed effects on the relevant response variable (e.g., time to capture, success rate, movement speed). Random effects were also incorporated into the models to account for the hierarchical and non-independent structure in the data at the level of individual animals. Model fitting was conducted using restricted maximum likelihood (REML) estimation, which provides unbiased estimates of variance and covariance parameters. We then used post hoc emmeans testing to obtain estimates of the fixed effects and assess potential significant influences of each variable and their interaction effects on the response variable.

To test whether mice avoided backtracking to recently visited objects during search, we classified each three-object sequence (triplet) traversed by the mouse as either X-Y-X, in which the mouse returns to the previously visited object, or X-Y-Z, in which it visits three distinct objects. Observed triplet probabilities were compared to a shuffled null, and per-triplet z-scores were computed for the search and pursuit states separately. We fit a linear mixed-effects model (z ~ triplet class × state + (1|animal/day), and used the X-Y-X vs. X-Y-Z class contrast within each state to test for a directional bias. The same model was used to assess whether the bias changed across training days. To test triplet class contrast in agent-based modeling data, we again used linear mixed-effects (z ~ triplet class × state) with a random intercept for run number.

To test whether object-directed behaviors changed across learning, we classified each object-directed frame during the search state into one of three behaviors per object and compared the proportion of each between the first and last days of object experience (day 1 vs. day 8). For each behavior we fit a binomial generalized linear mixed model (cbind(behavior, other) ~ day + (1 |animal/day)), with p-values FDR-corrected (Benjamini–Hochberg) across the nine behaviors. The same procedure was applied to the pursuit state.

Data are reported as mean *±* SEM unless otherwise noted. All tests were performed with a significance threshold of p < 0.05.

#### Object-occupancy divergence

To quantify how closely predator search occupancy matched the prey’s spatial distribution, we computed a total divergence from prey. For a given behavioral state (pursuit or search), we calculated the proportion of time the mouse spent nearest each object, along with the corresponding proportions for the roach, assigning each frame to its nearest object among the four objects (platform, ping-pong balls, long tube, and short tube). The total divergence was defined as the sum, across objects, of the absolute difference between the mouse and roach occupancy proportions:

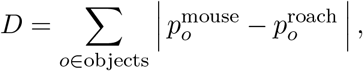

where 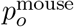 _and 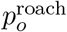_ denote the proportion of time the mouse and roach, respectively, spent nearest object *o*. Larger values indicate greater mismatch between predator search allocation and prey occupancy. For the experimental data, we computed *D* per animal per day and tested whether it differed between the first and last days of object exposure (day 1 vs. day 8) using a linear mixed-effects model (total divergence ~ day + (1 | animal)), fit separately for the pursuit and search states, with individual animal as a random effect. For the agent-based model (Fig. 5E), the same metric was computed from simulated occupancy proportions and compared between the value-matched and value-mismatched conditions with an analogous model (total divergence ~ condition + (1 | replicate)), with simulation replicate as a random effect.

#### Hyperparameter sweep

To assess whether the model’s key behavioral signatures depended on the specific parameter values used above, we performed a grid sweep over seven parameters governing the prey decision policy and predator search memory. We swept five values of each parameter, yielding 5^7^ = 78,125 combinations. The swept parameters were: `prey_top_n`, the number of top-scoring candidate flee moves the prey sampled among (the |*A*_top_| retained candidates in the prey policy); `prey_temperature`, the softmax temperature *T* controlling how uniformly the prey sampled among those candidates; `sterm_mem_min` and `sterm_mem_max`, the search-memory cooldown floor and ceiling (*T*_min_ and *T*_max_); `sterm_mem_tau`, the timescale over which the cooldown grew from floor to ceiling (*τ*_mem_); `prey_hide_bonus`, the concealment bonus added to a candidate flee move’s score for leading toward a concealment-capable tile (*β*); and `pexpire`, the per-turn probability that a hidden prey spontaneously became visible again. The single-value settings used elsewhere in the paper fall within these swept ranges.

For each parameter combination we computed two summary metrics. The *Mismatch*→*Match Realignment Score* quantified the benefit of a correctly informed search prior: it was the divergence between roach and mouse object-tile occupancy proportions during search under a value-mismatched (uninformed) predator, minus the same divergence under a value-matched (correctly informed) predator, so that higher values indicate greater improvement in occupancy alignment when the predator’s search prior matched the prey’s true hiding distribution (Fig. S5). The *Search*−*Pursuit Path-Style Contrast* quantified the expected behavioral split between states: it was the net path measure (loop count minus back-and-forth count) during search minus the same measure during pursuit, restricted to the value-matched condition, so that higher values indicate a combined preference for looping trajectories during search and direct back-and-forth trajectories during pursuit (Fig. S6).

To visualize the dependence of each metric on individual parameters, we summarized the sweep pairwise: each off-diagonal cell shows the metric averaged over all combinations sharing a given pair of row and column parameter values, marginalizing over the remaining five parameters, and each diagonal panel shows the full marginal distribution of the metric across all combinations for each value of a single parameter. Both signatures were robust across the swept parameter space.

## Data Availability

The datasets required to reproduce the main figures in this paper are provided as source data files. Raw video recordings are available from the corresponding author upon reasonable request, due to their large size.

## Code Availability

All relevant code to reproduce figures in this paper are publicly available at github.com/hengenlab/Schneider_McGregor

## Acknowledgments

This work is supported by National Institutes of Health (F31NS134240) and Wu Tsai Institute Computational Fellowship (AMS), Cure Alzheimer’s Fund (JNM, KBH), National Institutes of Health BRAIN Initiative (R01EB029852) (ELD), and National Institutes of Health (R01NS118442 and R01AG091773) (KBH).

We thank Kiran Bhaskaran-Nair for assistance with video processing and Kylee Kest for assistance with running experiments.

## Author Contributions

A.M.S. designed experiments, assisted in recordings, performed machine-learning, behavioral modeling, produced figures, and wrote the manuscript. J.N.M. performed statistical analysis, wrote the manuscript, and produced figures. M.S. performed data preprocessing, statistical analysis, and machine-learning. J.A. led the recordings. S.Z. performed data preprocessing, machine-learning, and behavior labeling. D.W. performed data preprocessing, machine-learning, and behavior labeling. J.T. performed data preprocessing and machine-learning. G.Y., E.E., J.C. and J.P. performed behavior labeling. V.S. assisted in recordings. E.L.D. provided mentorship and guidance on machine-learning. D.L. provided mentorship and guidance on behavioral modeling. K.B.H. contributed to initial concept, provided mentorship and guidance on recordings, and wrote the manuscript.

